# Establishment of a Long-Term Germ-Free Medaka Model Reveals Microbiota-Dependent Regulation of Growth, Immunity, and Metabolism

**DOI:** 10.64898/2026.03.09.710661

**Authors:** Pan-Pan Jia, Ming-Fei Wu, Li-Ping Ma, Feng-Yi Guo, Lan-Chen Zhang, Yan Li, De-Sheng Pei

**Author notes:** **Corresponding author:** E-mail address (D. S. P). These authors contributed equally to this work (Pan-Pan Jia, Ming-Fei Wu).

## Abstract

Germ-free (GF) animal models are indispensable for dissecting host-microbiota interactions and their roles in health and disease. The small teleost fish medaka (*Oryzias latipes*) provides unique advantages for establishing GF models across developmental stages, yet the functions of its intestinal microbiota and metabolites remain poorly characterized. Here, we developed both early-life and chronic GF medaka models to systematically characterize host biology in the absence of microbiota and evaluate the contribution of gut-derived metabolites to growth and immune development. Using a refined sterile feeding and verification protocol, we successfully maintained GF medaka for up to 57 days post-fertilization (dpf). As anticipated, GF fish displayed developmental delays, impaired organogenesis, reduced immune competence, and metabolic dysregulation. Supplementation with sterile gut-derived metabolites partially alleviated these deficits, as evidenced by enhanced locomotor activity and immune responses. Mechanistically, recovery was associated with improved ribosome biogenesis, tricarboxylic acid cycle activity, and histidine and pyruvate metabolism, suggesting enhanced protein synthesis and immune maturation. However, metabolite supplementation also elevated oxidative stress and inflammatory responses and failed to fully restore long-term survival or organ development. Our findings support the use of GF medaka as a versatile platform for investigating microbiota-host interactions across life stages. By integrating metabolite interventions, this model provides critical insights into the functional roles of gut microbiota and offers a valuable tool for advancing microbiome research in health and disease.

## Introduction

The gut microbiota exerts profound influence on host physiology, shaping immune development, metabolic balance, and neural function ^1–3^. Perturbations in microbial communities have been increasingly associated with chronic inflammatory, metabolic, and neurodevelopmental disorders. Germ-free (GF) animal models, which eliminate microbial confounders, have therefore become indispensable for establishing causal relationships between microbes, their metabolites, and host biology ^4–6^. Yet, despite their value, current GF systems remain constrained by limited scalability, species diversity, and applicability across lifespans.

Aquatic vertebrates have emerged as powerful complements to mammalian GF models, offering tractable genetics, optical transparency, and high-throughput potential ^6–8^. Zebrafish (*Danio rerio*) have been widely used to study microbial regulation of physiology, including probiotic-pathogen interactions and microbiota–mediated modulation of neuro-developmental pathways ^9–12^. GF systems have also been extended to rainbow trout (*Oncorhynchus mykiss*) and Atlantic cod (*Gadus morhua*) ^13,14^. However, most studies remain confined to short larval stages, typically within 10 days post–fertilization, leaving the biology of long–term or adult GF fish largely unexplored. The freshwater medaka, *Oryzias latipes* (*O. latipes*), offers distinct advantages for addressing these limitations, particularly for studies on sex-specific effects, as its XY sex-determination system mirrors that of humans and provides a closer parallel than zebrafish ^15^. *O. latipes* further combines high fecundity, daily spawning, and transparent embryos with a longer incubation period and earlier onset of feeding compared to zebrafish, enhancing larval survival and facilitating sterile rearing. These traits, together with their established use in developmental biology, genetics, and toxicology, position medaka as a promising yet underutilized GF model ^16,17^.

In parallel with advances in GF husbandry, attention has turned to gut–derived metabolites as key mediators of host-microbe interactions ^1^. Microbial metabolites, including bile acid derivatives, short–chain fatty acids, and amino acid catabolites, enter systemic circulation and modulate host physiology ^18^. Microbial bile salt hydrolase influences bile acid signaling and cholesterol metabolism ^19^, while short–chain fatty acids regulate energy balance and immune tolerance ^20^. In zebrafish, microbial metabolites have been shown to reverse neural gene expression deficits and enhance lipid absorption ^21^. Despite these insights, their roles in medaka remain largely unexplored.

Two major gaps remain unaddressed: the need for long-term GF medaka models and a systematic evaluation of microbial metabolites in this context. Here, we establish both short– and long–term GF medaka, extending survival into early adulthood, and investigate whether gut–derived metabolites can mitigate GF–associated phenotypes. This dual approach advances GF fish husbandry and provides mechanistic insights into how microbial metabolites influence immunity, metabolism, and development. Beyond gnotobiology, the work offers a scalable vertebrate platform for microbiome research with implications for precision medicine, aquaculture, and environmental health.

## Results

### Absence of microbiota induced growth retardation and behavioral suppression in *O.latipes*

The surface of fertilized medaka eggs (*O.latipes* and *O.melastigma*) is covered with villi and fibers that can harbor microorganisms. To establish GF models, fibers were minimized and eggs separated into individuals (**Fig. 1A**). To enhance sterility and hatching rates, embryos were cultured in 48-well plates, then transferred post-hatching to progressively larger vessels (24-well, 12-well, T25, and T75 flasks; **Fig. 1B**). Continuous microbial monitoring of feed and media using TSA and TSB under aerobic and anaerobic conditions confirmed sterility (**Fig. S1**). Hatching rates were significantly higher in GF O. latipes (OL-GF, 84.47%) than in GF marine medaka (MM-GF, 51.39%) (**Fig. S2**), while survival at 28 dpf was also greater in OL-GF (28.89%) than in MM-GF (21.53%), leading to the selection of O. latipes for subsequent experiments. GF medaka were established at 7, 14, 21, and 28 dpf, and comparison with CR controls revealed marked morphological divergence by 21 dpf (**Fig. 1C**). GF fish exhibited growth retardation, as evidenced by reduced survival, body length, body weight, and heart rate (**Fig. 1D-G**). Behavioral assays indicated a preference for static positioning and reduced activity (**Fig. 1H**, **Fig. S3A**), with significant decreases in distance, velocity, activity frequency, cumulative active duration, and manic behaviors (**Fig. 1I**, **J**; **Fig. S3D-G**), alongside elevated static frequency and duration (**Fig. 1K**, **L**).

**Fig. 1.**
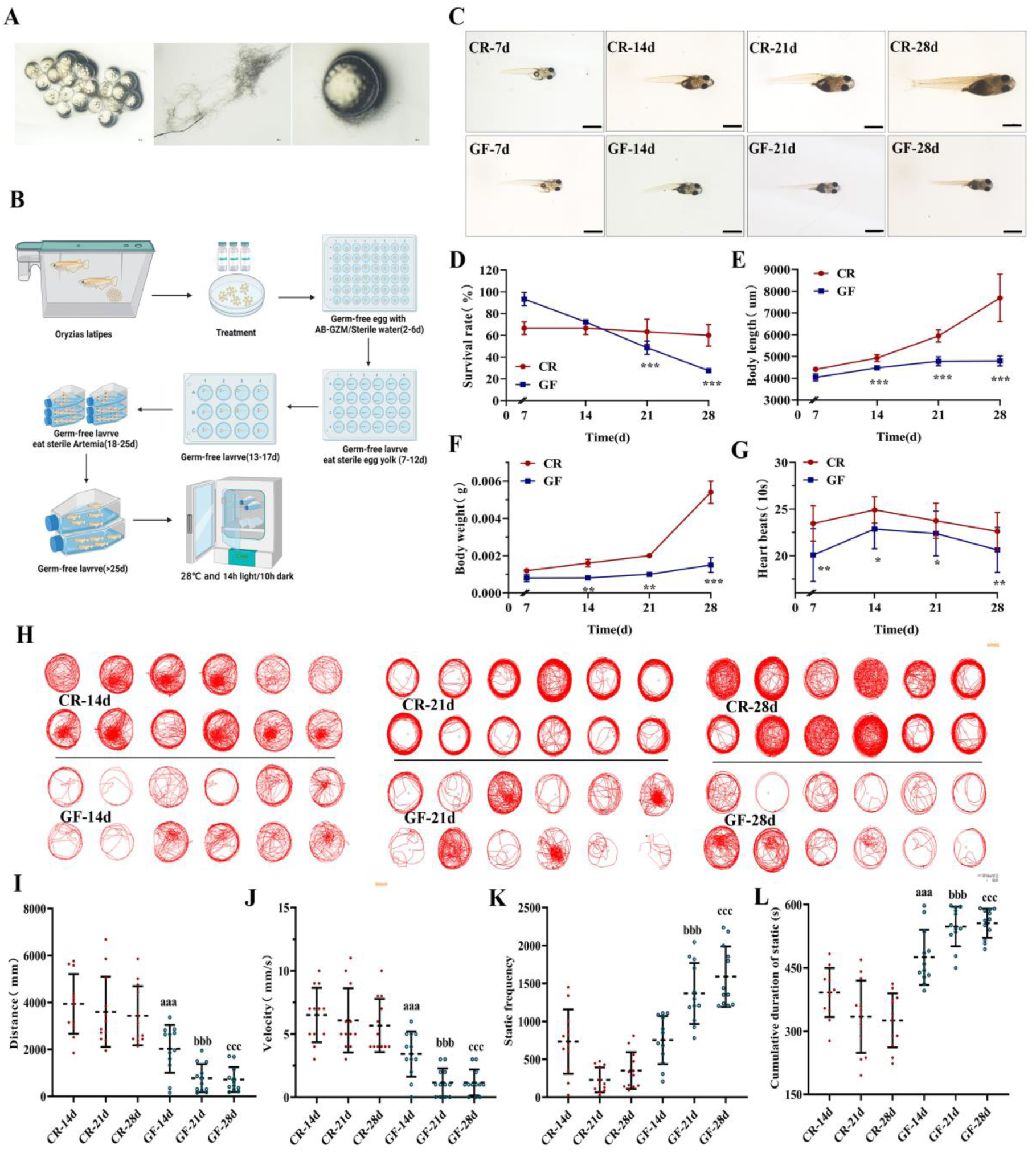
Long-term absence of microbiota induces growth retardation and behavioral suppression in GF *O.latipes*. (A) Fertilized eggs of *O.latipes*. (B) Flowchart of GF O. latipes model generation. (C) Morphology. Magnification: 1×; scale bar: 1000 μm. (D-G) Survival (%), body length, weight, and heartbeats (per 10 s).*, **, ***: GF vs. CR, P < 0.05, P < 0.01, P < 0.001. (H) Behavioral trajectory map. (I-L) Distance, velocity, static frequency, and accumulated static duration. aaa: GF–14 vs. CR–14, P < 0.001; bbb: GF–21 vs. CR–21, P < 0.001; ccc: GF–28 vs. CR–28, P < 0.001.

### Absence of microbiota induced low immunity during *O.latipes* growth

Histological analysis revealed distinct developmental patterns in GF *O.latipes*. By 28 dpf, both GF and CR fish exhibited fully developed major organs, though GF individuals showed delayed organogenesis (**Fig. 2A**). The GF liver displayed marked retardation, with hepatocyte hypertrophy, disrupted lobular architecture, and progressive cytopathology including vacuolation and necrosis, most evident at 21 and 28 dpf (**Fig. 2B**). Splenic development was also delayed, with loose parenchymal organization but preserved red/white pulp differentiation (**Fig. 2C**). Intestinal morphology showed reduced mucosal fold height, muscularis thickness, and villus width, along with increased absorptive vacuoles in GF-21d/28d groups (**Fig. 2D**). AB-PAS staining further indicated glycogen/neutral mucins in GF-28d intestines, contrasting with mixed mucins in CR-28d fish (**Fig. 2E**). To assess immune ontogeny, gene expression and cellular dynamics were profiled during larval to juvenile transition (**Fig. 3A-H**). In the CR fish, *bf/c2*, *c3*, and *lyz* expression increased with age, while *il-10*, *il-1β*, and *il-21* rose significantly at 14 d, and *ik* remained stable. GF groups showed suppressed *bf/c2*, *c3*, and *lyz* but elevated *il-10*, *il-21*, *il-6*, and *il-1β* relative to CR controls. Neutral red staining revealed progressive macrophage expansion in CR fish, whereas GF groups exhibited attenuated increases from 14 d onward, with 28 d counts significantly lower than CR (**Fig. 3I-L**). Sudan Black B staining showed neutrophil numbers peaking at 14 d in CR before declining, while GF groups displayed no significant stage-dependent changes (**Fig. 3M, N**).

**Fig. 2.**
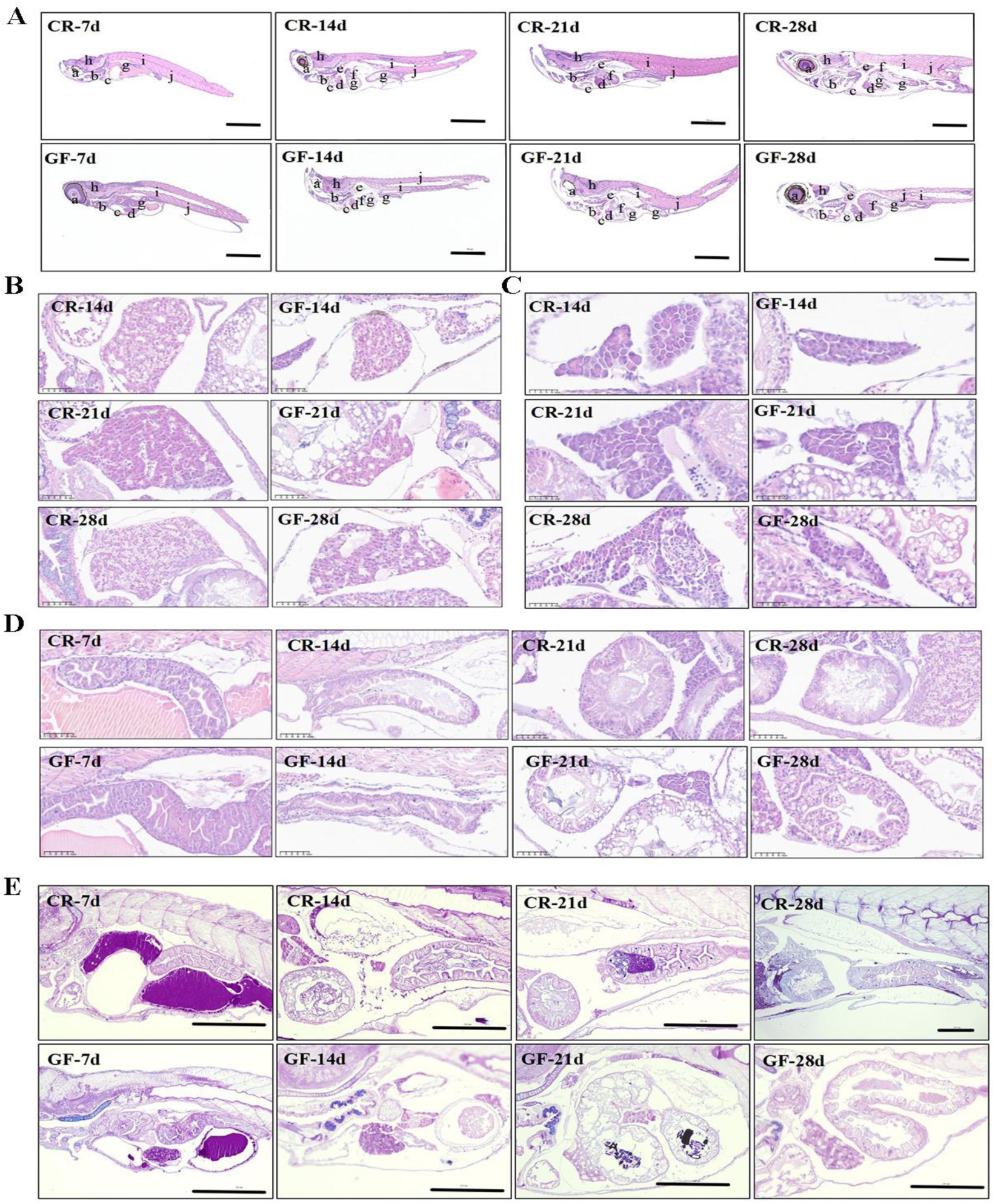
Analysis of tissue development in various organs of *O.latipes*. (A) Tissue development of multiple organs: a, eyes; b, gills; c, heart; d, liver; e, kidney; f, spleen; g, intestine; h, brain; i, backbone; j, muscle. Magnification: 4×; scale bar: 500 μm. (B-C) Liver and spleen development in GF O. latipes. Magnification: 80×; scale bar: 25 μm. (D) H&E staining of intestinal tissue. Magnification: 80×; scale bar: 25 μm. (E) AB-PAS staining of the intestine. Magnification of CR–28 d group: 10×; scale bar: 200 μm. Magnification of others: 20×; scale bar: 200 μm.

**Fig. 3.**
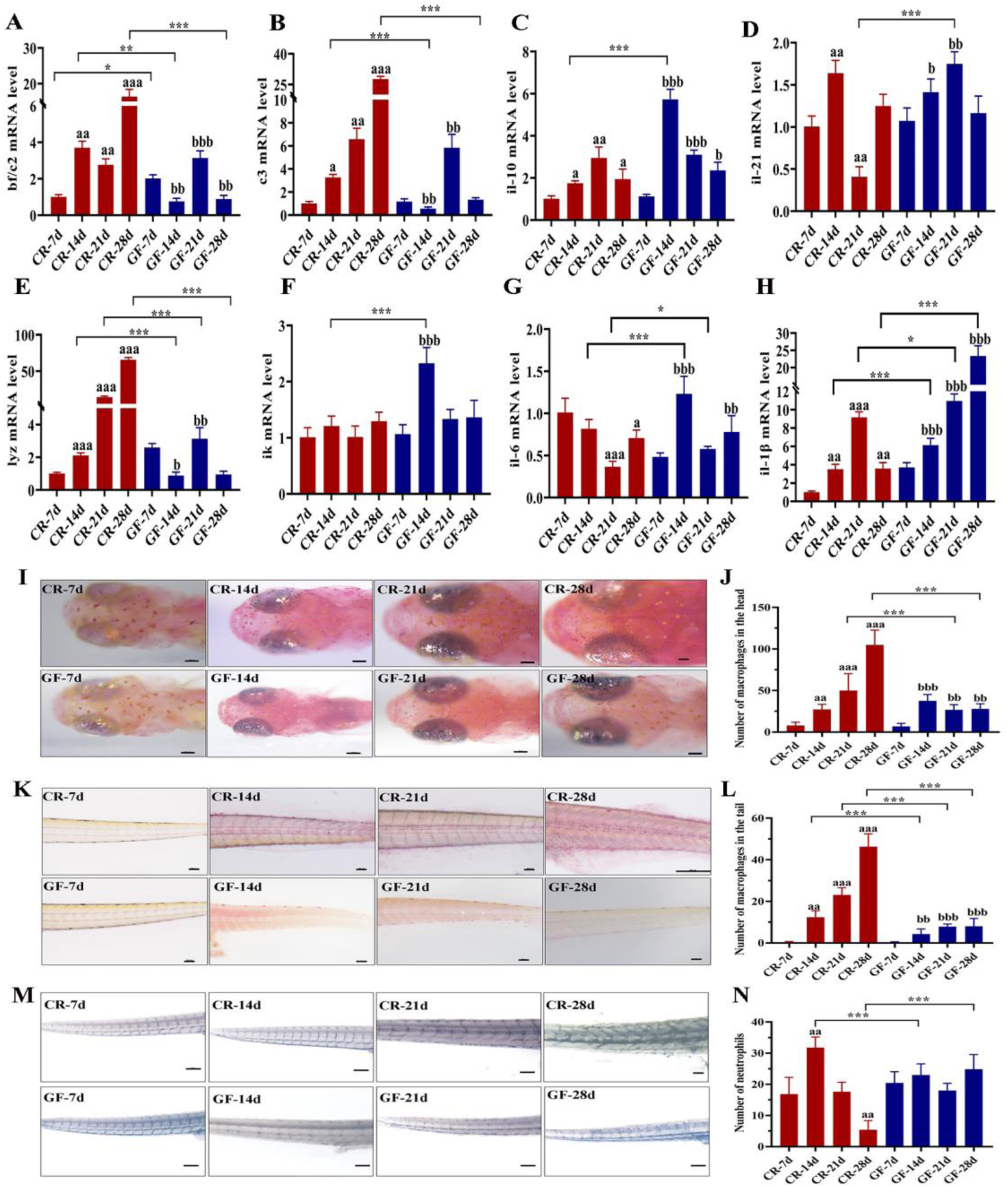
Long-term sterile environment induces low immunity in *O.latipes*. (A-H) Expression of *bf/c2*, *c3*, *il-10*, *il-21*, *lyz*, *ik*, *il-6*, and *il-1β* genes. (I-J) Macrophages in the head region. Magnification of CR–28 d: 6×; scale bar: 100 μm. Magnification of others: 8×; scale bar: 100 μm. (K-L) Macrophages in the tail region. Magnification of CR–28 d: 4×; scale bar: 100 μm. Magnification of others: 6×; scale bar: 100 μm. (M-N) Neutrophils. Magnification of CR–28 d: 4×; scale bar: 100 μm. Magnification of others: 6×; scale bar: 100 μm. a, aa, aaa: compared with CR–7 d, P < 0.05, P < 0.01, P < 0.001; b, bb, bbb: compared with GF–7 d, P < 0.05, P < 0.01, P < 0.001;*, **, ***: GF vs. CR, P < 0.05, P < 0.01, P < 0.001.

### Impacts of microbiota on gene expression profiles in CR and GF medaka

Transcriptomic analysis identified 1,627 differentially expressed genes (DEGs) between GF and CR groups, including 527 upregulated and 1,100 downregulated (**Fig. S4A**, **Fig. 4A**). GO enrichment highlighted extracellular region, extracellular matrix, and collagen-containing matrix components, with molecular functions related to heme binding, serine-type endopeptidase inhibition, and ECM structural constituents, and biological processes involving protein hydrolysis, immune response, and visual perception (**Fig. 4B**). KEGG analysis revealed enrichment in cytoskeletal organization, cytokine-cytokine receptor interaction, ECM-receptor interaction, cofactor biosynthesis, and phagosome pathways (**Fig. 4C**). qRT-PCR validation of immune-and phagosome-related genes confirmed RNA-seq reliability (**Fig. S4B-E**). Notably, immune-related genes, including complement components (*c8a*, *c8b*, and *c9*) and cytokine-associated genes (*cxcr4*, *cd4-1*, *il-1*, and *xcr1a.1*), were significantly downregulated in GF fish. Metabolomic profiling identified ten dominant metabolites: hypoxanthine, oleamide, L-phenylalanine, L-tyrosine, creatine, DL-tryptophan, N-indole-3-acrylic acid, L-methionine, 2-hydroxycinnamic acid, and L-isoleucine, accounting for ≥50% of total metabolites in both groups (**Fig. 4D-E**). In total, 471 metabolites differed between GF and CR fish, with 243 upregulated and 174 downregulated in GF (**Fig. S5A-B**). GF fish showed elevated palmitoylcarnitine, LPG 22:5, Lysopc 16:2 (2N isomer), and LPS 22:6, but reduced 4-guanidinobutanoic acid, trigonelline, 4-acetyl-4-(ethoxycarbonyl)heptanedioic acid, inosine 5′-monophosphate, and methandrostenolone (**Fig. S5C-D**). KEGG pathway analysis of these metabolites revealed enrichment in nucleotide, glutathione, and purine metabolism, as well as pantothenate/CoA biosynthesis, arginine/proline metabolism, and β-alanine metabolism (**Fig. 4F**).

**Fig. 4.**
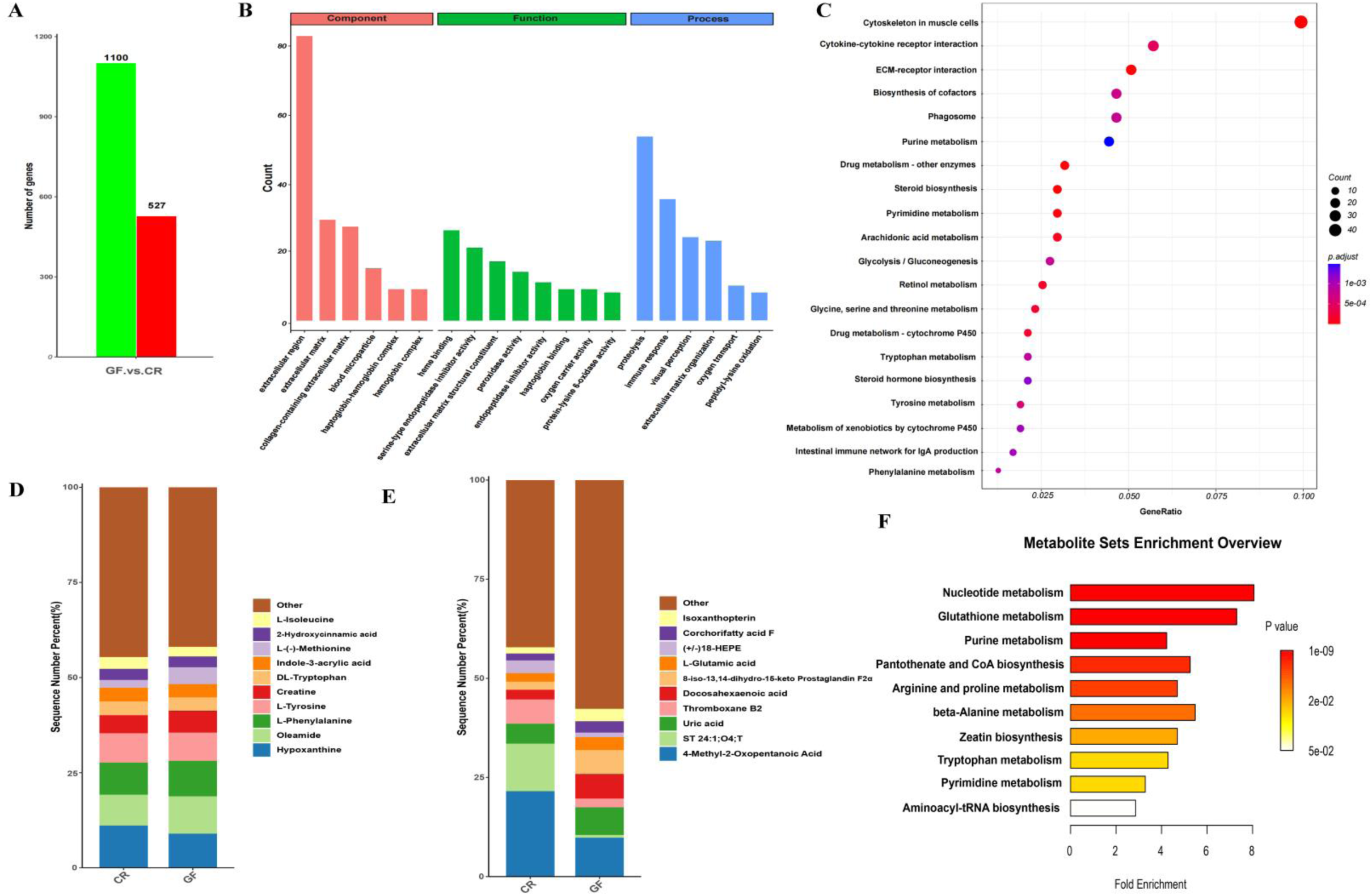
Lack of microbiota alters immune and metabolic gene expression profiles. (A) Bar chart of differentially expressed genes. (B) GO enrichment analysis of differentially expressed genes. (C) KEGG enrichment analysis of differentially expressed genes. (D-E) Percentage of metabolites in positive and negative ion modes. (F) KEGG enrichment map of differential metabolites.

### Intestinal metabolites partially restore locomotor and immune functions in GF *O.latipes*

To evaluate the effects of sterile gut-derived metabolites (M) on GF *O. latipes*, the optimal concentration was first determined, with 1×10⁻³ improving 7–day survival and selected for subsequent experiments (**Fig. S6A**). Compared with GF controls, the M-treated group showed higher survival, body length, body weight, and heart rate, though differences were not significant (**Fig. 5A-D**). Behavioral tracking revealed thigmotaxis (wall-hugging behavior) in both groups, with GF fish displaying more erratic trajectories (**Fig. 5E**). M treatment enhanced locomotor activity, increasing distance, velocity, activity frequency, cumulative active duration, and manic duration (**Fig. 5G-I**), while significantly reducing static frequency and duration (**Fig. S6B-D**). Flow cytometry showed GF medaka had elevated lymphoid cells and reduced granulocytes relative to CR controls, with no change in precursors. M treatment partially reversed this phenotype by decreasing lymphoid cells and increasing granulocytes, with a non-significant trend toward higher precursor levels (**Fig. 5J**, **K**).

**Fig. 5.**
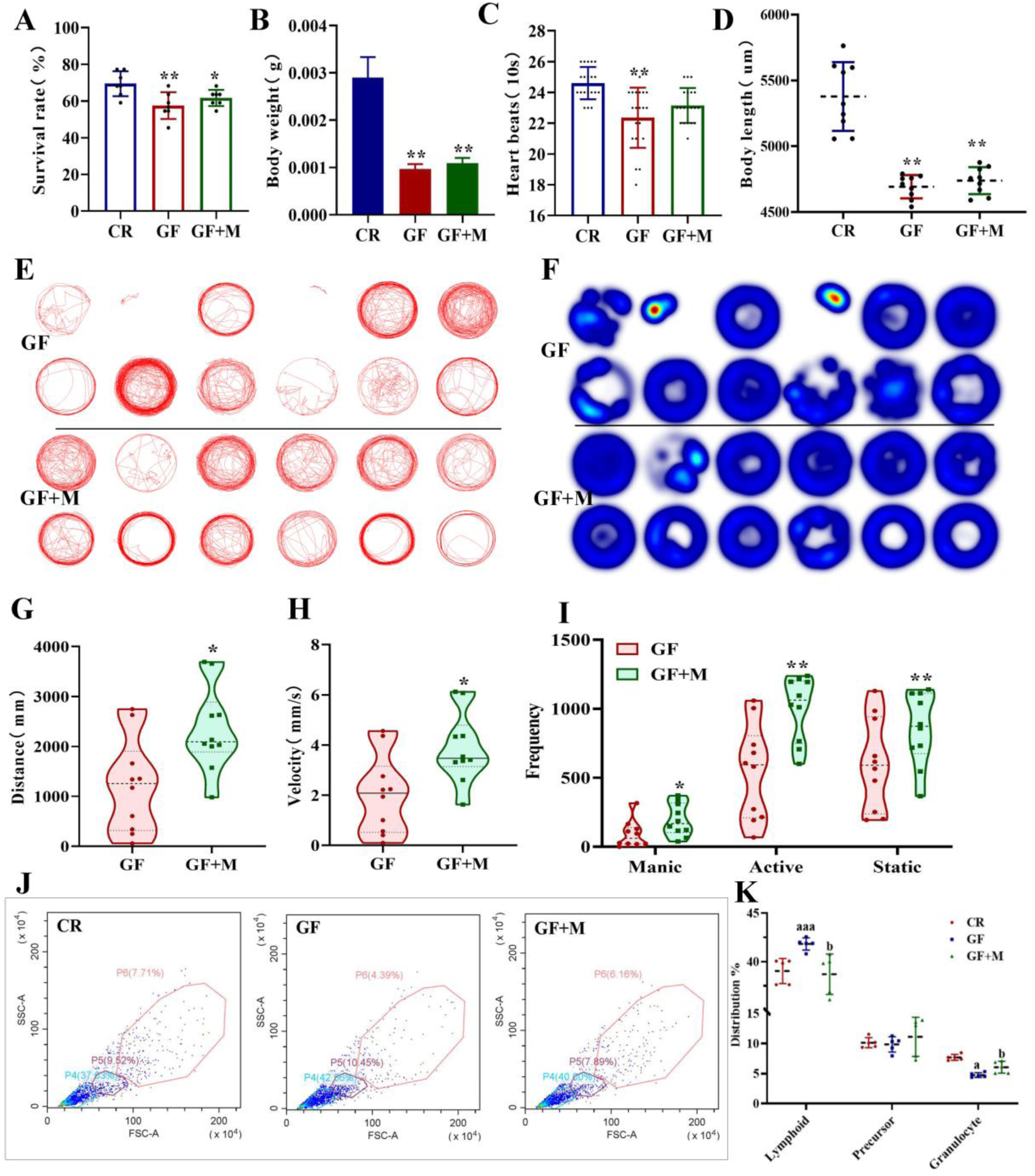
Gut-derived metabolites partially restore locomotor and immune functions in GF *O.latipes*. (A-D) Survival rate (%), body weight, heart rate (per 10 s), and body length of CR, GF, and GF+M groups.*, **, ***: GF and GF+M vs. CR, *P* < 0.05, *P* < 0.01, *P* < 0.001. (E) Behavioral trajectory map. (F) Behavioral heat map. (G-I) Distance, velocity, and frequency of manic, active, and static states. (J, K) Flow cytometry profiles of renal cells: P6, granulocytes; P5, precursors; P4, lymphocytes. a, aa, aaa: GF vs. CR, *P* < 0.05, *P* < 0.01, *P* < 0.001; b, bb, bbb: GF+M vs. GF, *P* < 0.05, *P* < 0.01, *P* < 0.001.

### Microbial metabolites demonstrated limited efficacy in rescuing long-term GF-induced developmental arrest in *O.latipes*

To assess the long-term biology of GF medaka, we examined survival and development. While CR medaka survival stabilized by 3 weeks, GF fish showed continuous decline with a maximum lifespan of 57 days, and gut-derived metabolites treated GF medaka (GF+M) displayed a similar pattern without lifespan extension (**Fig. 6A**). By 30 d, CR fish reached the late larval stage with fin development and pigmentation, and by 50 d attained the juvenile stage with mature morphology and pronounced pigmentation. In contrast, GF and GF+M groups exhibited incomplete scale formation and reduced dorsal fins (**Fig. 6B**), along with significantly reduced body length and weight; GF+M treatment showed only a non-significant trend toward improvement (**Fig. 6C**, **D**). Histology revealed complete maturation of digestive, respiratory, cardiovascular, and nervous systems in the CR-50d group, whereas the GF group displayed delayed organogenesis and pathological changes across liver, intestine, and muscle (**Fig. 6E**). CR livers were enlarged with compact hepatocytes and enhanced metabolic capacity, while GF livers were smaller with disorganized architecture (**Fig. 6F**). Intestinal analysis showed CR-50d fish had thickened mucosa, well-developed villi, and robust digestive function, whereas GF intestines exhibited thin mucosa, vacuolated enterocytes, underdeveloped villi, and impaired barrier function (**Fig. 6G**). AB-PAS staining confirmed abundant glycogen and acidic mucopolysaccharides in CR-50d intestines, while GF fish showed minimal acidic mucus secretion, with no improvement from M treatment (**Fig. 6H**).

**Fig. 6.**
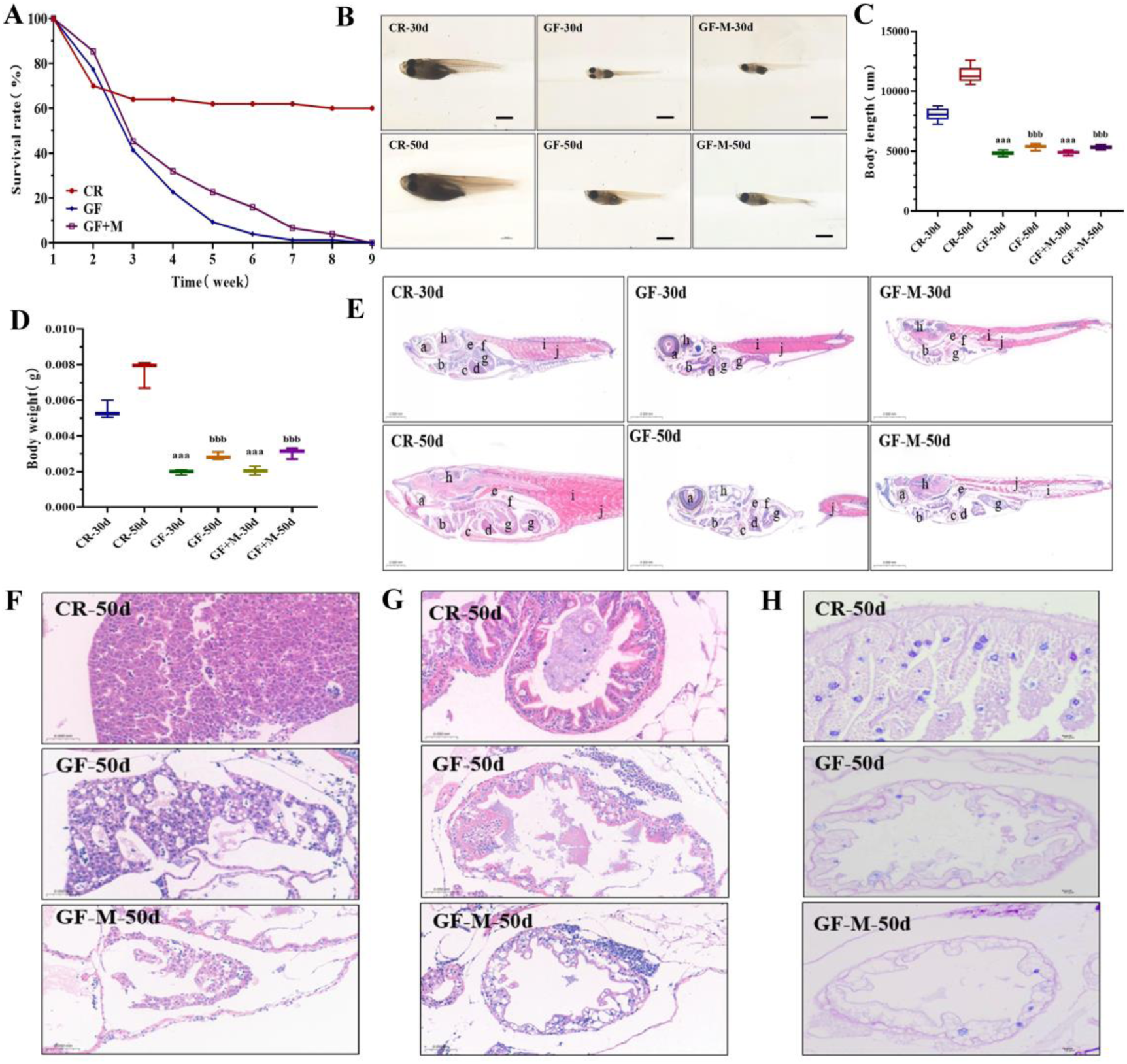
Gut microbial metabolites show limited efficacy in rescuing long-term GF-induced developmental arrest in *O.latipes*. (A) Long-term survival rate (%). (B) Morphology of CR and GF at 30 d and 50 d. Magnification of CR-50 d: 0.75×; scale bar: 1000 μm. Remaining images: 1×; scale bar: 1000 μm. (C, D) Body length and body weight of CR and GF at 30 d and 50 d.*, **, ***: GF vs. CR, *P* < 0.05, *P* < 0.01, *P* < 0.001. (E) Organs: a, eye; b, gills; c, heart; d, liver; e, kidney; f, spleen; g, intestine; h, brain; i, trunk; j, muscle. Magnification: 5×; scale bar: 500 μm. (F) Liver tissue development. Magnification: 40×; scale bar: 50 μm. (G, H) H&E staining of intestine. Magnification: 80×; scale bar: 25 μm. (I) AB-PAS staining of the intestine. Magnification of CR-28 d: 10×; scale bar: 200 μm. Magnification of others: 20×; scale bar: 200 μm.

### Gut microbial metabolites modulated translation and metabolism but failed to fully restore ECM and immune homeostasis in GF *O. latipes*

To elucidate the molecular effects of metabolite M in GF medaka, we performed integrated transcriptomic and metabolomic analyses. PCA distinguished the three experimental groups (**Fig. 7A**), and clustering showed that M treatment partially restored GF gene expression patterns (**Fig. 7B**). Compared to GF controls, the GF+M group exhibited 674 upregulated and 1,277 downregulated genes (**Fig. 7C**). Immune-related responses were evident: GF+M significantly increased *bf/c2*, *c3-1*, *c3-2*, and *lyz* expression but not *ik* (**Fig. S7A**), while *il-10* and *il-1β* were downregulated and *il-21* and *tnf-α* were upregulated (**Fig. S7B**).

**Fig. 7.**
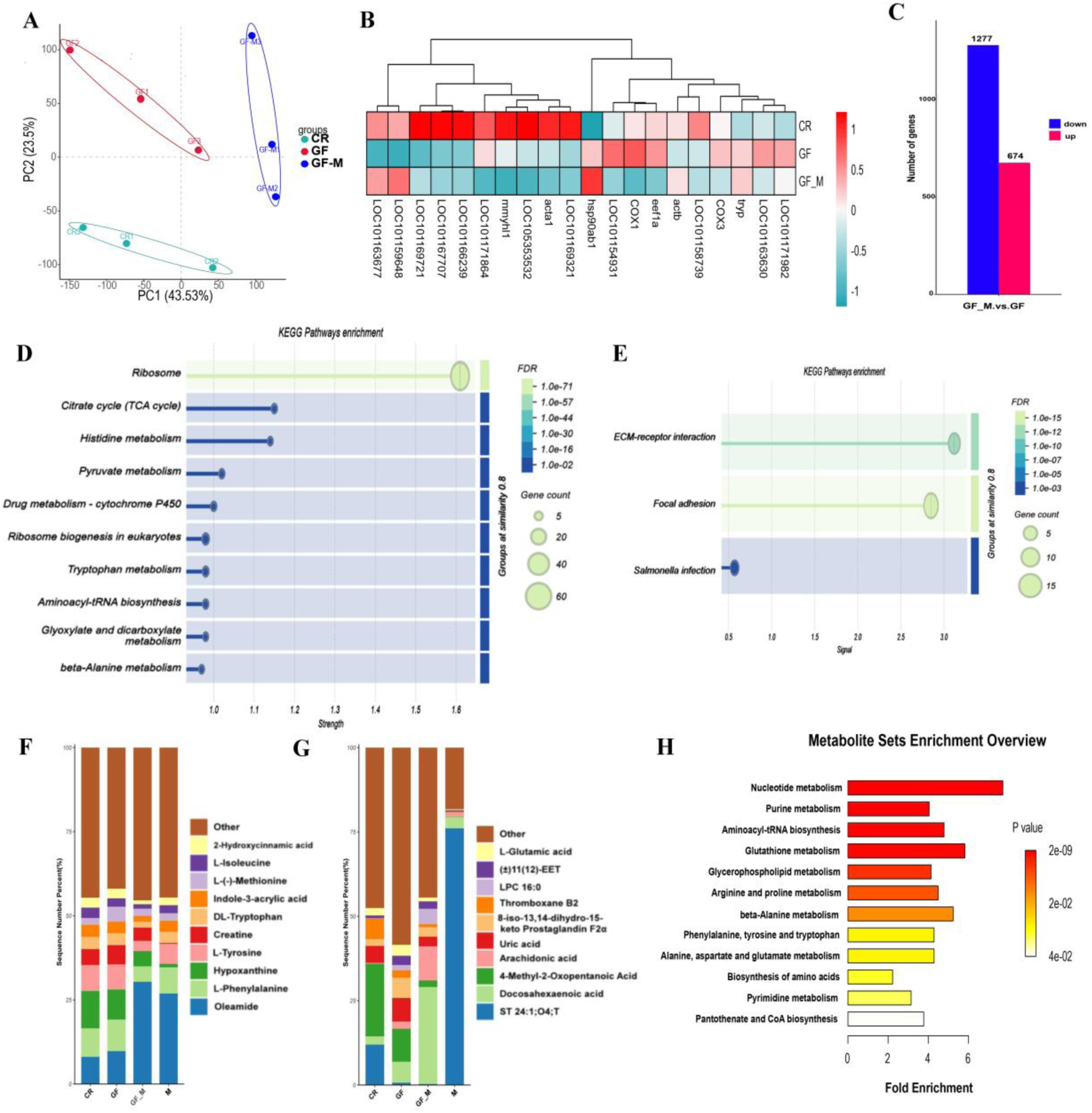
Gut microbial metabolites modulate translation and metabolism but fail to fully restore ECM and immune homeostasis in GF *O.latipes*. (A) Scatter plot of principal component analysis (PCA). (B) Cluster plot of differentially expressed genes. (C) Cluster and Venn plots of differentially expressed genes. (D-E) KEGG enrichment analyses of differentially expressed genes. (F-G) Percentage of metabolites in positive and negative ion modes. (H) KEGG enrichment map.

GO enrichment of rescued genes highlighted translation-related complexes, processes, and functions (**Fig. S8A-C**), and KEGG analysis revealed enrichment in ribosome biogenesis, the TCA cycle, and histidine/pyruvate metabolism (**Fig. 7D**), suggesting regulation of protein synthesis and energy metabolism. Persistently altered genes were enriched in extracellular matrix (ECM)-related categories (**Fig. S9A-C**), with KEGG pathways implicating ECM-receptor interaction and focal adhesion (**Fig. 7E**), indicating sustained ECM disruption and altered host-pathogen dynamics.

Metabolomic profiling identified oleamide, L-phenylalanine, hypoxanthine, L-tyrosine, creatine, and DL-tryptophan as dominant metabolites in positive ion mode (**Fig. 7F**), and ST 24:1; O4; T, docosahexaenoic acid, 4-methyl-2-oxopentanoic acid, arachidonic acid, and uric acid in negative ion mode (**Fig. 7G**). In total, 245 metabolites differed between GF and GF+M groups, with 71 upregulated and 36 downregulated (**Fig. S10A-B**). Box plots highlighted increases in PC O-20:3, LPE 20:0, and CAR 20:4 (positive mode) and LPE 17:2, LPC 22:6, and LPS 22:6 (negative mode), alongside decreases in QLH, tyrosylalanine, and ethyl-β-D-glucuronide (**Fig. S10C-D**). KEGG annotation emphasized nucleotide, purine, and glutathione metabolism as the most affected pathways (**Fig. 7H**).

## Discussion

This study successfully established a sustainable GF medaka *O.latipes* model, enabling long-term maintenance from embryogenesis through a lifespan of up to 57 days. By addressing the technical challenges posed by adhesive egg surface structures and optimizing sterilization, hatching, and feeding protocols, we achieved complete sterility while maintaining a hatchability rate of 84.47%, which exceeded that of conventional systems (70.27%). This improvement parallels findings in GF rainbow trout ^14^, likely reflecting a reduction in microbial infection. The methodological advances described here extend previous work in GF zebrafish and trout ^14,22^, underscoring the feasibility of medaka as a tractable vertebrate model for host-microbiota research.

Food autoclaving induces significant nutrient loss in fish feed ^23^. To address this, we used live GF *Artemia nauplii* in later stages, which are characterized by high nutritional value, including abundant protein and unsaturated fatty acids. In accordance with the developmental needs of *O. latipes*, a multi–phase feeding protocol was implemented. Newly hatched larvae were provided with sterile egg yolk for the first 3-5 days. From days 6-10, the diet was supplemented with powdered *Artemia*. Between days 11-20, a transition to a combination of powdered and live *Artemia* took place. Beyond this period, feeding consisted exclusively of live *Artemia*. Despite these technical achievements, GF medaka exhibited profound developmental, behavioral, and physiological impairments. Growth retardation became evident after 14 days post-hatching, accompanied by hypoactive locomotor behavior and abnormal social interactions. Histopathological analyses confirmed delayed fin formation, impaired organogenesis, and severe intestinal underdevelopment, resulting in nutritional deficiencies and reduced survival. These phenotypes mirror those reported in GF mammals, reinforcing the evolutionarily conserved role of microbiota in vertebrate growth, organ maturation, and energy homeostasis ^24–26^. Notably, the maximum lifespan of GF medaka (57 days) exceeded that of previously reported GF zebrafish, suggesting species-specific differences in resilience to microbiota deprivation ^27,28^.

At the immunological level, GF medaka displayed marked deficiencies in innate immune development. Expression of complement components (c3, c2) and lysozyme was significantly reduced, while macrophage and neutrophil populations failed to expand appropriately. These findings are consistent with the pivotal role of gut microbiota in regulating early immune competence in teleosts ^29–31^. Integrated transcriptomic and metabolomic analyses further revealed broad downregulation of immune- and growth-related pathways, coupled with depletion of immunomodulatory metabolites, such as adenine, estrone, and LPC (13:1-sn1). Together, these results highlight the indispensable role of microbiota in sustaining mucosal barrier integrity, metabolic regulation, and immune homeostasis^32–34^. The observed metabolic disturbances, including impaired nutrient utilization, reduced short-chain fatty acid (SCFA) production, and consequent malnutrition, likely exacerbate immunodeficiency19.

Notably, supplementation with sterile intestinal metabolites partially rescued several GF phenotypes. Behavioral activity, complement expression, granulocyte counts, and protein synthesis capacity were improved, suggesting that microbial metabolites act as key mediators of host development and immune regulation ^19^. Transcriptomic signatures indicated enhanced ribosomal biogenesis, chaperone activity, and metabolic pathways, while metabolomic profiling revealed restoration of essential fatty acids such as docosahexaenoic acid (DHA) and arachidonic acid (ARA), both critical for membrane integrity and anti-inflammatory signaling ^20^. However, extracellular matrix (ECM)-related pathways remained uncorrected, and persistent risks of oxidative stress and inflammation were evident. These findings underscore both the therapeutic potential and the limitations of metabolite-based interventions, highlighting the complexity of host-microbiota crosstalk.

The multifaceted effects of microbial metabolites on cytokine regulation further highlight their systemic influence ^20,35^. For example, GF+M medaka exhibited suppression of *il-10* alongside increased *tnf-α* and *il-21*, while reducing *il-1β*. IL-21, in particular, is a key cytokine that promotes B-cell proliferation, antibody production, and T-cell differentiation^36,37^. Moreover, SCFAs and indole derivatives produced by commensals, such as *Lactobacillus*, modulate mucosal homeostasis and gut barrier integrity ^1,38^. These findings align with evidence that microbial metabolites exert influences beyond the gut, modulating immune responses in distal organs, including the central nervous system ^39^.

## Conclusions

This study establishes a long-term GF *O.latipes* model, sustaining sterility from embryos to 57 days and revealing hallmark consequences of microbiota absence, including developmental delay, organ immaturity, immunodeficiency, and metabolic dysregulation. Supplementation with intestinal-derived metabolites partially restored locomotor activity, immune responses, and protein synthesis, though with potential risks of oxidative stress and inflammation. The microbial controllability and genetic tractability of medaka uniquely position this system to dissect microbiota-dependent regulation of immunity, metabolism, and development, while offering translational value for modeling immune-mediated disorders such as inflammatory bowel disease and for testing interventions ranging from defined metabolite supplementation to probiotics or fecal microbiota transplantation. Future efforts should prioritize systematic screening of key microbial metabolites and the rational design of functional microbial consortia to optimize host health and identify novel therapeutic strategies.

## Materials and Methods

### Ethics statement

All fish experiments were conducted in accordance with the Guide for the Care and Use of Laboratory Animals (8th Edition, 2011, ILARCLS, National Research Council, Washington, D.C.) and the Chinese national standard for laboratory animal welfare (GB/T 35892-2018).

### Experimental fish and rearing conditions

To compare to our previous study of GF marine medaka *Oryzias melastigma* (*O.melastigma*) ^40^, fertilized eggs were collected from both wild-type *O.melastigma* and *O.latipes* under strict institutional animal care guidelines. *O. latipes* were maintained in a recirculating system with purified water, while *O. melastigma* were reared in artificial seawater at 35‰ salinity. Both systems were kept at 28 ± 1 °C under a 14 h light/10 h dark cycle, and larvae were fed freshly hatched brine shrimp nauplii (*Artemia* sp.) twice daily. Medaka reached sexual maturity at approximately 3 months of age and were paired at a female-to-male ratio of 1:2. Spawning typically occurred during the first hour of the light cycle, with each female producing 10-30 eggs suspended by chorionic filaments at the cloacal opening. Collected eggs were separated in Petri dishes after removal of nonviable eggs and debris, then incubated in methylene blue-supplemented embryo medium at 28 °C with daily water changes and removal of dead eggs. Hatching occurred at approximately 7 days post-fertilization (dpf) for *O. latipes* and 14 dpf for *O. melastigma*.

### Establishment and identification of GF *O.latipes* models from larvae to early adult stages

Fertilized eggs with normal development were selected under a stereomicroscope and sterilized in a biosafety cabinet using a protocol modified from standardized GF fish models ^12^. Eggs were washed three times with sterile GZM solution, three times with sterile AB-GZM solution, and incubated in sterile AB-GEM solution at 28 °C for 12-24 h. Sterilization included treatment with 0.4 g/L povidone-iodine (PVP-I) for 1 min followed by three GZM washes, immersion in 0.02 g/L sodium hypochlorite (NaClO) for 15-30 min with three final GZM washes, and transfer to 48-well plates containing AB-GZM until hatching. After hatching, larvae were maintained in 24-well plates and received daily sterile feeding and medium replacement. They were transferred to new vessels every 5-7 days. Feeding was divided into four phases: sterile egg yolk for 3-5 days, egg yolk plus sterile powdered Artemia nauplii for 6-10 days, powdered and live sterile Artemia for 11-20 days, and exclusively live sterile Artemia thereafter. Sterility was regularly monitored, and contaminated eggs were removed. Detailed protocols for reagent preparation, feed processing, *O.melastigma* egg sterilization, and sterility identification are provided in **Text S1**, with sterility assessment based on our previous study ^12^.

### Growth and developmental indicators of CR and GF medaka

The hatching rate, survival rate, body length, body weight, and heart rate of both conventionally raised (CR) and GF medaka were recorded at different developmental stages. Morphological comparisons were performed using a stereomicroscope, with photographs taken for documentation. Body length was measured as the linear distance from the snout to the caudal fin tip using the microscope’s integrated measurement tool. Hatching rate was calculated as (hatched larvae/viable embryos) × 100%, with embryos showing fully ruptured chorions considered hatched. Survival rate was calculated as (surviving larvae/initial embryos) × 100%, with embryos exhibiting whitening, absence of heartbeat, or loss of vital signs classified as deceased. Sterility rate was calculated as (confirmed GF medaka/total surviving GF medaka) × 100%. For each treatment group, 10-20 medaka were randomly selected for analysis, and all experiments included three biological replicates.

### Behavioral ability of CR and GF *O.latipes* models

Behavioral performance of CR and GF *O.latipes* at 14, 21, and 28 dpf was assessed using the DanioVision™ system, with 12 or 24 randomly selected individuals per group monitored by high–speed infrared cameras. Before testing, fish were acclimated for 10 min in 12– or 24–well plates containing test solution (one fish per well), maintained at 28 ± 1 °C. A 10-minute free–swimming session was recorded at 25 frames per second under controlled quiet conditions, and video data were analyzed using EthoVision XT® software.

### Histopathological analysis of tissue and organ development

Three to five medaka from each group were anesthetized with 100 mg/L MS-222 and fixed in 4% paraformaldehyde for at least 24 h. After ethanol dehydration and EDTA decalcification, samples were embedded in paraffin and sectioned at 5 µm using a microtome. Sections were mounted on glass slides, deparaffinized, and stained with hematoxylin-eosin (H&E) and Alcian blue-periodic acid-Schiff (AB-PAS). Following dehydration and clearing, slides were coverslipped for permanent preservation. All sections were digitally scanned and analyzed for organ development using K-Viewer software.

### Detection of immune cells *in vivo* by neutral red and sudan black B staining

Neutral red, a common dye for staining live fish, is taken up by phagocytic cells such as macrophages *via* lysosomes. Medaka (n = 6 per group, three replicates) were placed in 6–well plates and incubated with 10 μg/mL neutral red solution at 28 °C for 6-12 h in the dark, after which macrophage numbers in the head and tail regions were recorded by stereomicroscopy. Sudan Black B staining was used to label committed hematopoietic neutrophils. Embryos were first immersed in 0.003% 1–phenyl–2–thiourea, and larvae at various stages were fixed overnight in 4% paraformaldehyde at 4 °C. After washing with 1× PBST, samples were stained with Sudan Black B for 1 h on a shaker and destained with 70% ethanol until normal coloration was restored. The staining solution was prepared by mixing 30 mL of Solution A (0.6 g Sudan Black B in 200 mL absolute ethanol) with 20 mL of Solution B (16 g phenol in 30 mL absolute ethanol plus 100 mL of 30 g/L disodium hydrogen phosphate), followed by filtration. Six medaka from each group were analyzed, and the procedure was repeated three times.

### Expression of key genes related to immune in CR and GF medaka

Based on medaka size, 3-15 individuals from each group were collected in triplicate. Total RNA was extracted using TRIzol reagent after homogenization, and genomic DNA was removed by DNase treatment (details in **Table S1**). cDNA was synthesized using the reverse transcription system described in **Table S2** under the following conditions: 50 °C for 15 min, 85 °C for 5 s, then held at 4 °C. Quantitative PCR was performed with the reaction system in **Table S3** using the following thermal cycling parameters: 95 °C for 3 min, followed by 40 cycles of 95 °C for 10 s, 60 °C for 30 s, and 72 °C for 30 s. Primer sequences were designed using the NCBI-Primer tool and are listed in **Table S4**. Relative gene expression was calculated using the 2–ΔΔCT method with β–actin as the housekeeping gene.

### Transcriptome Sequencing Analysis

Total RNA was extracted from 10 medaka larvae at 21 dpf per group (three biological replicates) using the TRIzol method. RNA quality was assessed by agarose gel electrophoresis, NanoPhotometer spectrophotometry, and an Agilent 2100 Bioanalyzer to confirm purity, concentration, and integrity, with three technical replicates per sample. Libraries were constructed and subjected to high–throughput sequencing, and differentially expressed genes were identified and analyzed using Gene Ontology (GO) and Kyoto Encyclopedia of Genes and Genomes (KEGG) enrichment. Detailed methodologies are provided in **Text S2**. The raw transcriptome sequencing data are accessible at the NCBI SRA under BioProject ID PRJNA1404584.

### Untargeted metabolomic profiling

For metabolomic analysis, 100 medaka larvae per group (three biological replicates) were ground in liquid nitrogen and extracted with 500 µL of 80% methanol. After centrifugation, the supernatant was diluted with mass spectrometry–grade water to a final methanol concentration of 53%, recentrifuged, and subjected to LC–MS analysis (detailed in **Text S3**). Metabolites were annotated using the KEGG (https://www.genome.jp/kegg/pathway.html), HMDB (https://hmdb.ca/metabolites), and LIPIDMaps (http://www.lipidmaps.org/) databases. The metabolomics data have been deposited to the MetaboLights database (https://www.ebi.ac.uk/metabolights/) under accession number MTBLS13709.

### Preparation and exposure of gut-derived metabolites from *O.latipes*

Following the methods of Rea et al. ^41^, intestinal metabolites were extracted from adult *O.latipes* (n=5). After euthanasia by ice bath for ≥10 min, intestinal tissues were isolated, placed in pre–chilled 1× PBS, and homogenized at a 1:3 (w/v) ratio by vortexing for 1 min to release luminal contents. The suspension was centrifuged at 14,000 × g for 30 min at 4 °C, filtered through a 0.22 μm membrane for sterilization, and stored at −20 °C. For exposure experiments, metabolites were diluted (1:100-1:1000) to simulate physiological levels in medaka intestines. Briefly, 30 mL of sterile GZM medium was added to 25 cm² culture flasks containing five larvae, and thawed metabolite stock was introduced at graded concentrations (1:100, 1:500, 1:1000), with 200 μL approximating the metabolites from 2.7 larval intestines. During exposure, culture medium was regularly sampled for sterility testing using the methods described in **Text S1**.

### Flow cytometric analysis of immune cells in *O.latipes*

Single-cell suspensions were prepared from the head kidney (primary hematopoietic organ) and spleen of *O.latipes* (10 fish per group, three replicates). Fresh tissues were transferred to ice-cold PBS (1 mL with 1% FBS) and mechanically dissociated through a 40 μm cell strainer. Primary cells were treated with 3-5 volumes of pre-chilled erythrocyte lysis buffer, gently vortexed, and incubated at room temperature for 90 s, followed by centrifugation (500 × g, 4 °C, 5 min) to remove lysed erythrocytes. Cells were washed twice with ice-cold PBS (1% FBS) by centrifugation (500 × g, 4 °C, 3 min) to ensure purity. Before analysis, cells were stained with 2 nM propidium iodide (PI) for 5 min to exclude dead cells (PI-negative). Immunophenotyping was performed on a BD FACSLyric flow cytometer, with lymphocyte, monocyte, and granulocyte subpopulations distinguished by forward scatter (FSC-A) and side scatter (SSC-A). Data were analyzed using FlowJo v10.6.2 (BD Biosciences) with gating strategies optimized by isotype and fluorescence-minus-one (FMO) controls.

### Statistical analysis

Statistical analyses were performed using SPSS version 20.0. Student’s t-test was applied for two-group comparisons, while one-way ANOVA was used for multiple-group analyses. Pairwise comparisons were conducted using the LSD and Dunnett’s methods. Data are presented as mean ± standard deviation (SD) of group replicates, with significance defined at P < 0.05, 0.01, or 0.001 compared with control groups.

## Supporting information

https://identifiers.org/ncbi/insdc.sra:PRJNA1404584

## Acknowledgments

We would like to extend our sincere gratitude to the anonymous reviewers for their time, effort, and valuable insights. Their constructive comments and suggestions significantly improved the quality of this manuscript.

## Author contributions

**P.P.J**. Validation, Formal analysis, Data curation, Writing-Original draft preparation;

**M.F.W.** Formal analysis, Data curation, Methodology, Writing-Original draft preparation; **L.C.Z.**, **F.Y.G.**, **L.P.M.,** and **Y.L.** Validation and Visualization. **D.S.P.** Conceptualization, Writing-Reviewing and Editing, Supervision, Project administration, Funding acquisition.

## Data availability statement

The authors confirm that all data supporting the findings of this study are included in the article and its supplementary materials. The “Transcriptome sequencing data” is available and can be accessed via the following DOI or accession ID: https://identifiers.org/ncbi/insdc.sra:PRJNA1404584; The “Metabolomic profiling data” is accessible through the DOI or accession ID: https://identifiers.org/ metabolights: MTBLS13709.

---

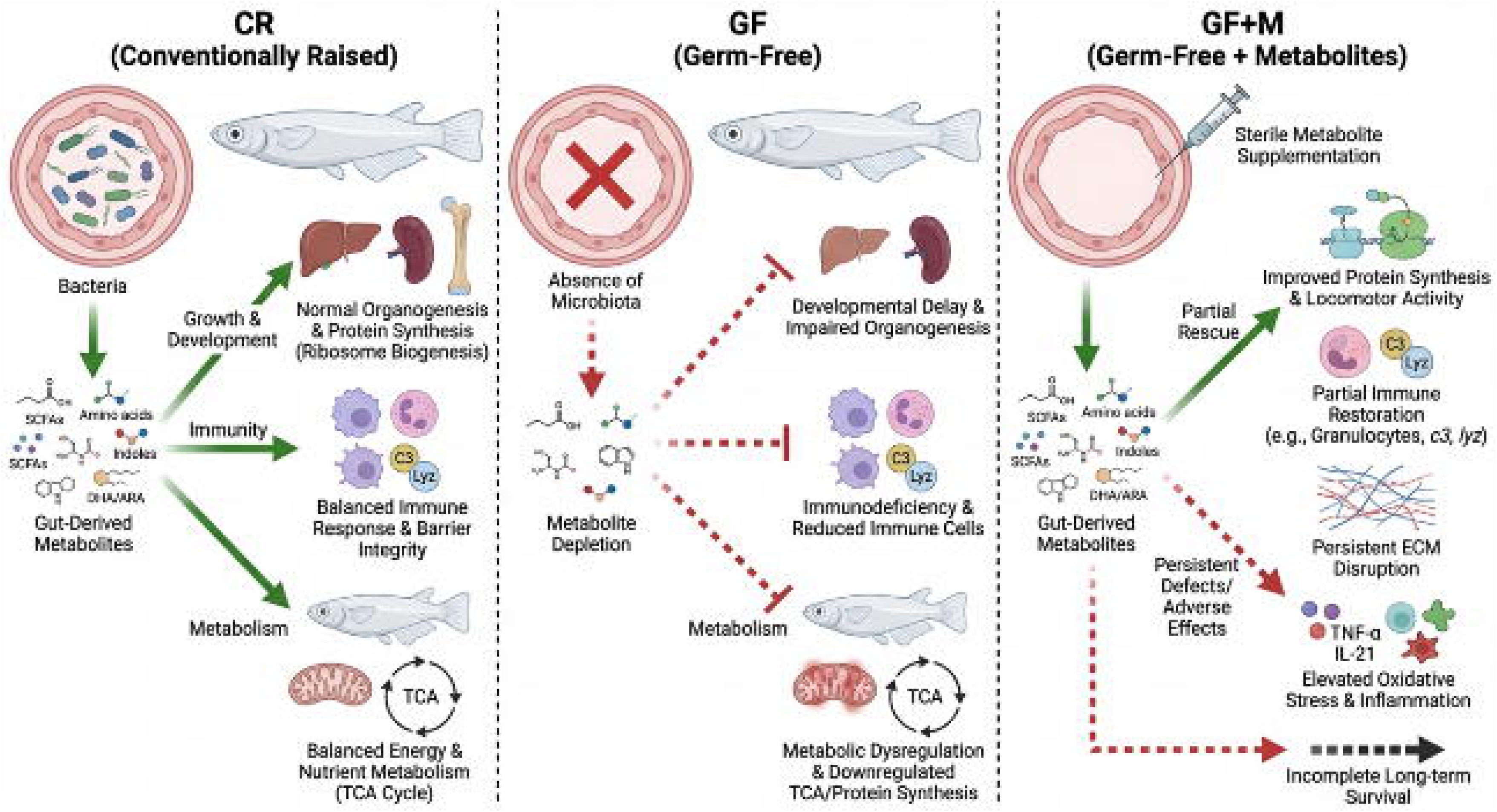

