## Supplementary material for "Establishment of a Long-Term Germ-Free Medaka Model Reveals Microbiota-Dependent Regulation of Growth, Immunity, and Metabolism": https://identifiers.org/ncbi/insdc.sra:PRJNA1404584

[*]**Corresponding author：**

### Text S1: Establishment of Germ-Free *Oryzias latipes* Model

#### 1. Sterile reagent preparation

The GZM solution was prepared by adding 600 μL of 4% sea salt solution (40 g/L) to 400 mL of sterilized water, followed by autoclaving. The AB-GZM solution was prepared by mixing 200 mL of GZM solution (made from 200 mL water and 300 μL of 4% sea salt solution), 400 μL of amphotericin B stock solution (250 μg/mL), 200 μL of kanamycin stock solution (10 mg/mL), 2 mL of ampicillin stock solution (20 mg/mL), and 400 μL of penicillin–streptomycin (100×), and then filter-sterilizing the mixture through a 0.22 μm membrane before storage at –20 °C. The final concentrations of antibiotics were: amphotericin B, 500 ng/mL; kanamycin, 10 μg/mL; ampicillin, 200 μg/mL; and penicillin–streptomycin, 20 U/mL. The procedure for sterilizing marine medaka fertilized eggs was similar to that for freshwater medaka, except that the GZM solution was replaced with MMEM (35‰ sterile artificial seawater). The MMEM solution was prepared by separately autoclaving 35 g of sea salt and 1 L of reverse osmosis (RO) water before mixing.

#### 2. Sterile feed preparation

For sterile egg yolk, hard-boiled egg yolk was suspended in sterile water by repeated pipetting, aliquoted into 1.5 mL microcentrifuge tubes, and homogenized using a tissue homogenizer to reduce particle size. The samples were sterilized twice by autoclaving at 120 °C, 103.4 kPa, for 30 min. The resulting sterile egg yolk was stored at –20 °C. For live sterile Artemia, 0.15 g of cysts were rinsed for 5 min in a sterile solution of 48 mL sterile RO water and 2 mL of 6% NaClO. The cysts were filtered through an 80-mesh sieve and washed three times with sterile RO water to remove residual NaClO. Sterilized cysts were transferred to a 100 mL wide-mouth bottle containing GF-Artemia hatching solution (1.5 g iodized salt in 60 mL RO water) and incubated on a shaker at 150 rpm and 28 °C under 2000 lux illumination for 36–48 h. After hatching, the mixture was left undisturbed for 1 min in a biosafety cabinet to allow separation of shells and larvae. Nauplii were collected from the bottom using a 3 mL Pasteur pipette and washed twice with sterile RO water to remove residual hatching solution. The harvested sterile Artemia were stored at 4 °C for up to 3 days. To facilitate ingestion by GF *Oryzias latipes* larvae, the Artemia were further homogenized using a tissue homogenizer and stored at –20 °C.

#### 3. Sterility identification

Feed samples were collected for microbial testing, including autoclaved egg yolk solution, live GF Artemia after two days of storage, and powdered live GF Artemia. Culture medium samples were also collected from the GF freshwater medaka system at 1, 5, 7, 10, and 14 days post-fertilization (including egg membranes and medium after dehulling), and every 5 days thereafter. For microbial examination, 0.2 mL of each sample from the sterile feed and rearing process was inoculated onto TSA solid and TSA anaerobic solid media, and into TSB liquid and TSB anaerobic liquid media. Solid media were incubated at 30 °C under aerobic or anaerobic conditions, while liquid cultures were shaken at 150 rpm and 30 °C. Cultures were monitored for 3–7 days, and microbial growth was recorded.

The TSA aerobic and anaerobic solid media contained 6 g tryptone, 2 g soytone, 2 g sodium chloride, and 6 g agar in 400 mL distilled water. After mixing and dissolving, the medium was sterilized by autoclaving at 120 °C for 30 min. Anaerobic media were purged with nitrogen before autoclaving. All media were poured into Petri dishes under sterile conditions (or in an anaerobic chamber for anaerobic variants) and stored at 4 °C.

The TSB aerobic and anaerobic liquid media were prepared with 6 g tryptone, 2 g soytone, and 2 g sodium chloride in 400 mL of distilled water. Once dissolved, the medium was aliquoted into anaerobic tubes if applicable, purged with nitrogen for anaerobic versions, sterilized at 120 °C for 30 min, and stored at 4 °C.

**Text S2: Transcriptome Sequencing Analysis**

During library construction, the NEBNext® Ultra™ RNA Library Prep Kit (Illumina) was used according to the manufacturer's instructions. Briefly, poly(A)+ mRNA was selected using oligo(dT) magnetic beads and randomly fragmented with divalent cations in NEB Fragmentation Buffer. The fragmented mRNA was used as a template to synthesize first-strand cDNA with random hexamer primers and M-MuLV Reverse Transcriptase. Second-strand cDNA was synthesized using DNA Polymerase I and RNase H. The double-stranded cDNA was purified, end-repaired, adenylated at the 3′end, and ligated to sequencing adapters. cDNA fragments of 250–300 bp were selected using AMPure XP beads, amplified by PCR, and purified again with AMPure XP beads to obtain the final library. The library was quantified with a Qubit 2.0 Fluorometer and diluted to 1.5 ng/μL. Insert size was assessed using an Agilent 2100 Bioanalyzer, and effective concentration was determined by qRT-PCR (⩾2 nM) to validate library quality. Libraries passing quality control were pooled according to effective concentration and desired sequencing depth. Sequencing was performed on an Illumina platform to generate 150 bp paired-end reads using Sequencing by Synthesis (SBS) technology. During sequencing, four fluorescently labeled dNTPs, DNA polymerase, and adapter primers were introduced into the flow cell for amplification. Each incorporation of a dNTP released a fluorescent signal, which was captured by the sequencer and converted into sequence reads by computational processing.

The high-throughput sequencer converted image data of sequenced fragments into sequence reads using the CASAVA pipeline, and the resulting data were stored in FASTQ format containing nucleotide sequences and corresponding quality scores. Raw reads often included adapter sequences or low-quality regions. To ensure data quality, raw reads were filtered to remove those containing adapter sequences, undetermined bases (N), or low-quality sequences (where >50% of bases had Qphred $\leq$ 20). After filtering, Q20, Q30, and GC content were calculated for the cleaned data. The reference genome and annotation files were downloaded from a genomic database. HISAT2 v2.0.5 was used to build a genome index and align paired-end clean reads to the reference genome. Novel transcripts were assembled and quantified using StringTie (v1.3.3b), which employs a network flow algorithm and de novo assembly. Gene expression levels were estimated as FPKM (fragments per kilobase of transcript per million mapped reads). For differential expression analysis between two groups (two biological replicates each), DESeq2 (v1.16.1) was applied. P-values were adjusted using the Benjamini–Hochberg method to control the false discovery rate, and genes with adjusted $P < 0.05$ were considered differentially expressed. In cases without biological replicates, edgeR was used. Read counts were normalized using the trimmed mean of M-values (TMM) method in edgeR, and differential expression was determined with adjusted P-values via the Benjamini–Hochberg procedure, using $|\log_2(\text{fold change})|$ and adjusted P as significance thresholds.

**Text S3: Untargeted Metabolomic Profiling**

A quality control (QC) sample was prepared by pooling equal volumes from each experimental sample. A blank sample was prepared using a 53% methanol–water solution, which underwent the same pretreatment procedure as the experimental samples. Chromatographic separation was performed on a Hypersil Gold C18 column maintained at 40 °C with a flow rate of 0.2 mL/min. For positive ion mode, mobile phase A consisted of 0.1% formic acid, and mobile phase B was methanol. For negative ion mode, mobile phase A was 5 mM ammonium acetate (pH 9.0) and mobile phase B was methanol. The mass spectrometry scan range was set from 100 to 1500 m/z. Electrospray ionization (ESI) source parameters were configured as follows: spray voltage 3.5 kV, sheath gas flow 35 psi, auxiliary gas flow 10 L/min, ion transfer tube temperature 320 °C, S-lens RF level 60, and auxiliary gas heater temperature 350 °C, in both positive and negative ion modes. Data-dependent acquisition (DDA) was applied for MS/MS scanning.

Raw data files were imported into the CD3.1 library search software for processing. Initial filtering of metabolites was based on retention time and mass-to-charge ratio. Peak alignment across samples was performed with a retention time tolerance of 0.2 min and a mass tolerance of 5 ppm to improve identification accuracy. Peak extraction was then carried out under the following criteria: mass tolerance 5 ppm, signal intensity deviation 30%, signal-to-noise ratio $\geqslant 3$, minimum signal intensity, and inclusion of adduct ions. Peak areas were quantitatively analyzed. Target ions were integrated, and molecular formulas were predicted using molecular and fragment ion information, followed by matching against the mzCloud (https://www.mzcloud.org/), mzVault, and Masslist databases. After removing background ions using blank samples, quantitative data were normalized using the formula: (sample quantitative value) ÷ (sum of metabolite values in the sample ÷ sum of metabolite values in the QC1 sample) to obtain relative peak areas. Finally, compounds with a coefficient of variation (CV) of relative peak area >30% in QC samples were excluded, yielding the final metabolite identification and relative quantification results.

**Table S1. The DNA enzyme reaction system (10 mL)**

| reagent | application amount (μL) |
|---|---|
| 5×gDNA Eraser Buffer | 2 |
| gDNAEraser | 1 |
| Total RNA | 1 |
| RNase Free $dH_2O$ | up to 10 μL |

**Table S2. Reverse transcription reaction system (10 μ L)**

| reagent | application amount（μL） |
|---|---|
| Remove the DNA enzyme reaction system | 10 |
| Prime Script RT Enzyme Mix Ⅰ | 1 |
| RTPrime Mix | 1 |
| 5×Prime Script Buffer | 4 |
| RNase Free $dH_2O$ | 4 |

**Table S3. RT-qPCR Reaction System (10 μL)**

| reagent | application amount（μL） |
|---|---|
| SYBR（TB-green II） | 5 |
| DEPC | 3.2 |
| 10 μM forward primer | 0.4 |
| 10 μM reverse primer | 0.4 |
| cDNA | 1 |

**Table S4. The primer sequences of *Oryzias latipes* models in this study.**

| Gene | Forward primer（5’to 3’） | Reverse primer（5' to 3'） |
|---|---|---|
| *β-actin* | CCACCATGTACCCTGGAATC | GCTGGAAGGTGGACAGAGAG |
| *c3* | CCTAAACAGCAAGCACAGACTCAC | CCCAGCATCAAAGAACACACT C |
| *bf/c2* | ATCGCCTTGGACATTTCAGAGA | GACACAGTGAAGGCTGCAATCT |
| *lyz* | ACCAATGCCATCAACCACAA | GTTGACTCTTCCGGTTTTTCC A |
| *il-21* | CTGCTGCAGCCTCAAAGCGT | TGA GGTGCAGTTTTGGCACGA |
| *ik* | TGACCAAGCAATCAACAGCG | GCTTGTGGAGGCCATACAAA |
| *il-1β* | GTCCAGCTGAACATGTCTAC | TTGTCTCCTTCTTGGTGGCA |
| *il-6* | CTGAAGCAGGTGGAGAAGGAGTAC | GACCCGCTCGCTCCTTTTCATCTT |
| *il-10* | CCATTAAGAGCGAGTTCGC | ATCCTGCCGCCGCTTTGGG |
| *xcr1a.1* | CCGAGCGGTCAAGTTGGTGTTC | GCAGCAGTGAGAGAAGGCGATG |
| *tnfb* | ATTTACCTGGGCGGCGTGTTTAG | CTCTGCTCGGATGTGGTCTTGTTC |
| *mstnb* | TAGCCGTCACCTCAGCAGAACC | CATCGCAATCCAAGCCAGAGTCTC |
| *inhbab* | ATGCCTCTCCAACTGCCTCCTC | CGCTTCACTGCTTCCACCATCTC |
| *il-1* | ATCAGAATGGCAGCAGCAAGGC | GGCACCAGCATCAACGAAGGAG |
| *cd4-1* | GTGGGCGTGGTTGGTCTTCATAG | GGGCTGAGGGAGAGGACAGTTAC |
| *gh1* | CTCCACCTGCTCGCCCAGAG | TGTGTGCCGTGCTTGTCTAATGG |
| *epor* | CAGGTAGAAGTGGCACGAGCAAG | TCACTCCAGGCACTCCAGTAGC |
| *cxcr4* | GGGCGTCATGGAAGAACTCAACTC | TCCCACCAGAAAGCAGAGAGACAG |
| *c9* | ATTCTCCCTTGGCTTTGGTGTTCC | CACGGCGAATCTGGCTGACTG |
| *c8b* | ATTCTCCCTTGGCTTTGGTGTTCC | CACGGCGAATCTGGCTGACTG |
| *c8a* | TGTGCCACCATCATGCCGATTC | GGCTCTCGCAGAACGCATCG |
| 101166121 | ATCGCTTTCGGGATCAGAGTGTTG | CCTGTGCCTGAACTGGTGAACC |
| 101163459 | GAACACTCTGATCTGCCACGTCTC | GATGCTTCCTGACACCTTCTCACC |
| 101175656 | TGTGCTGTTTAGGTCGCCCAATC | CGCTGCCAACTTTCTGCCAAATG |
| 101156150 | CTGGGTCTGGTGGTGGGTCTG | GGCCCGCCCGATGGTTTATTC |
| 101163710 | CTCTGCCTCCTCTTCCTCACTCTG | GCTGCTGAACCTGATGACCTCTTC |

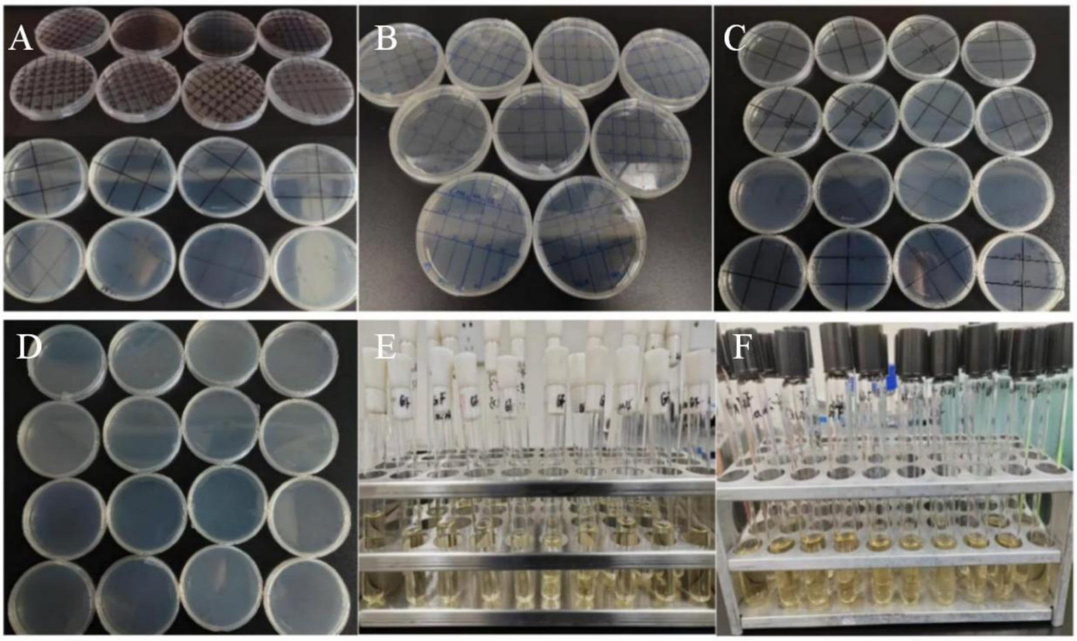


**Fig. S1 The detection of GF fish models with the samples of daily.**

(A-C) TSA aerobic solid medium; (D) TSA anaerobic solid medium; (E) TSB aerobic liquid medium; (F) TSB anaerobic liquid medium.

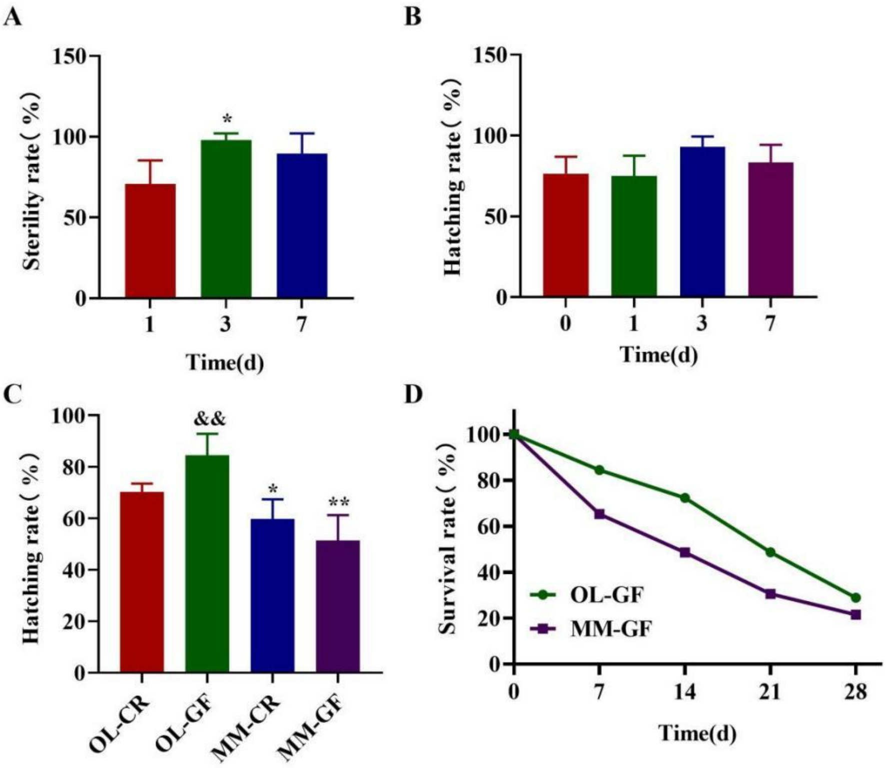


**Fig. S2. Development of GF *Oryzias latipes* and *Oryzias melastigma* models.** (A–B) Sterility rate (%) and hatching rate (%) using sterile AB-GZM reagent; *: 3 d vs. 1 d, $P < 0.05$. (C) Hatching rate of *O. latipes* and *O. melastigma* under common and sterile feeding conditions; *, and **: MM vs. OL, $P < 0.05$, $P < 0.01$; &&: OL-GF vs. OL-CR, $P < 0.01$. (D) Survival rate of GF *O. latipes* and *O. melastigma*.

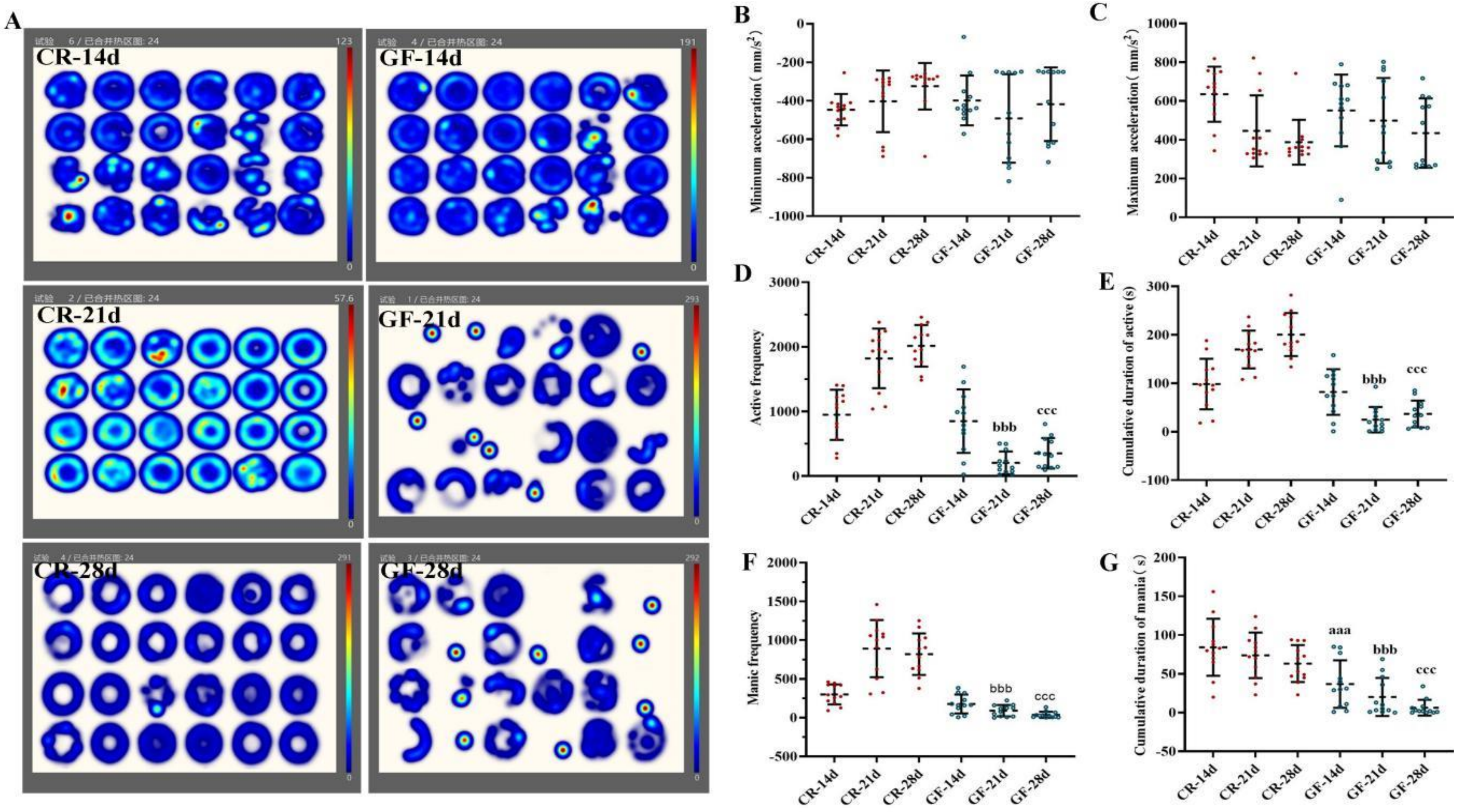


**Fig. S3. Behavioral analysis of GF *Oryzias latipes*.** (A) Behavioral heat map. (B–G) Minimum acceleration, maximum acceleration, accumulated active duration, active frequency, static frequency, and accumulated manic duration. aaa: GF-14 d vs. CR-14 d, $P < 0.001$; bbb: GF-21 d vs. CR-21 d, $P < 0.001$; ccc: GF-28 d vs. CR-28 d, $P < 0.001$.

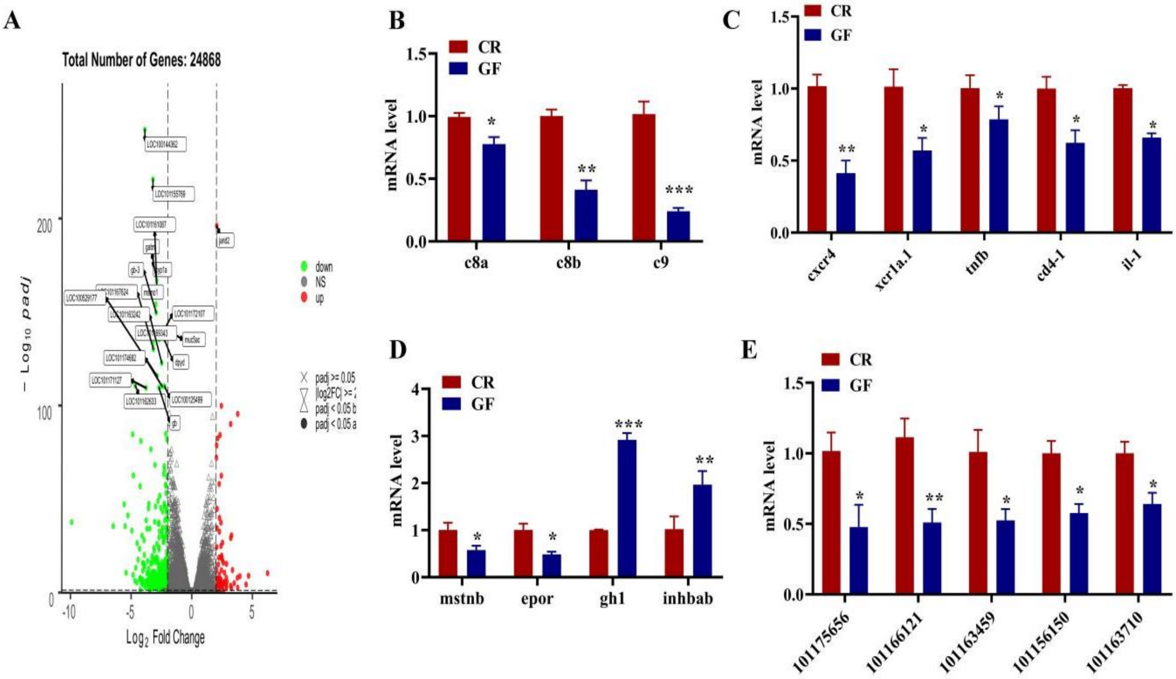


**Fig. S4. Transcriptome analysis of GF *Oryzias latipes*.** (A) Volcano plots of differentially expressed genes. (B–E) Validation of differentially expressed genes. *, **, and ***: GF vs. CR, $P < 0.05$, $P < 0.01$, $P < 0.001$.

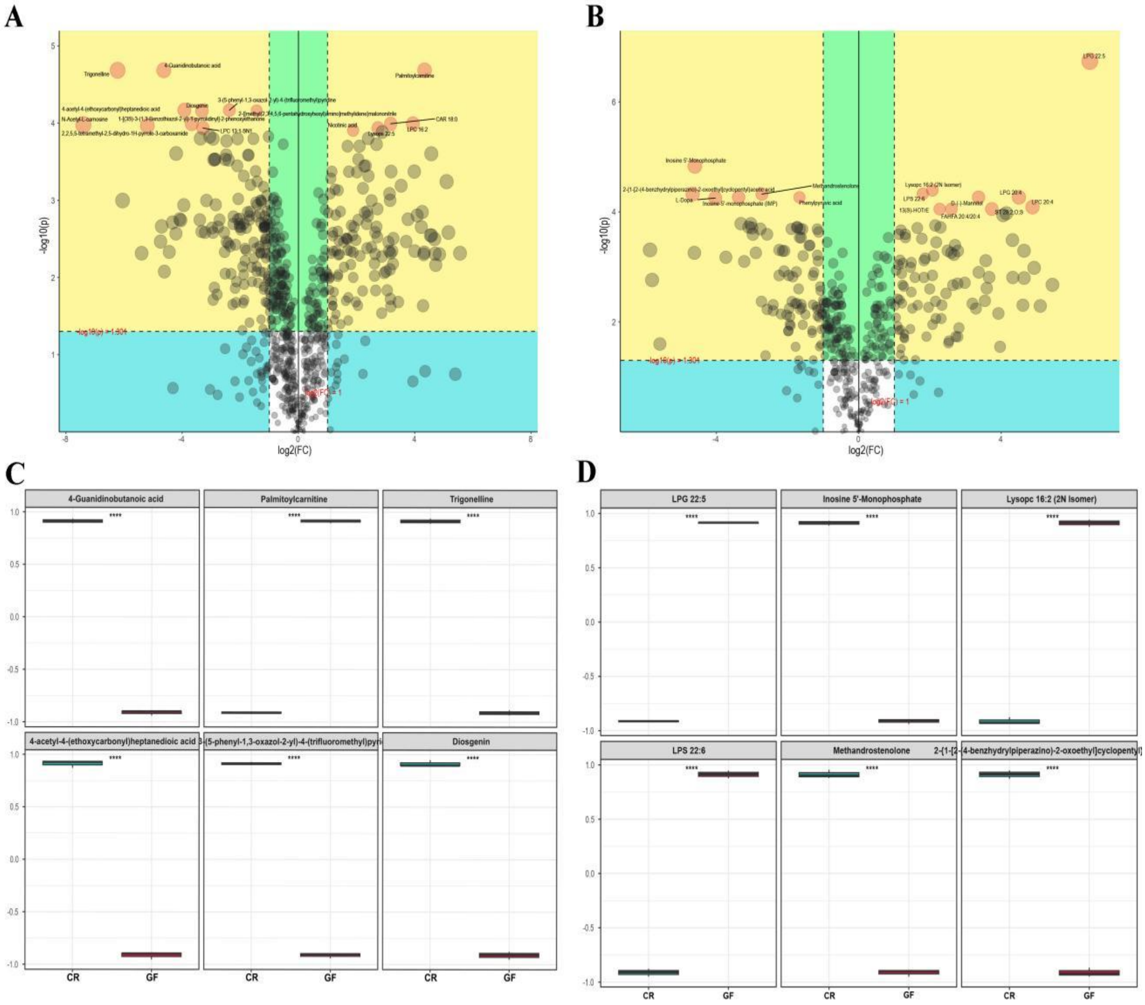


**Fig. S5. Metabolome analysis of GF *Oryzias latipes*.** (A–B) Volcano plots of differential metabolites in positive and negative ion modes. (C–D) Box plots of differential metabolites in positive and negative ion modes.

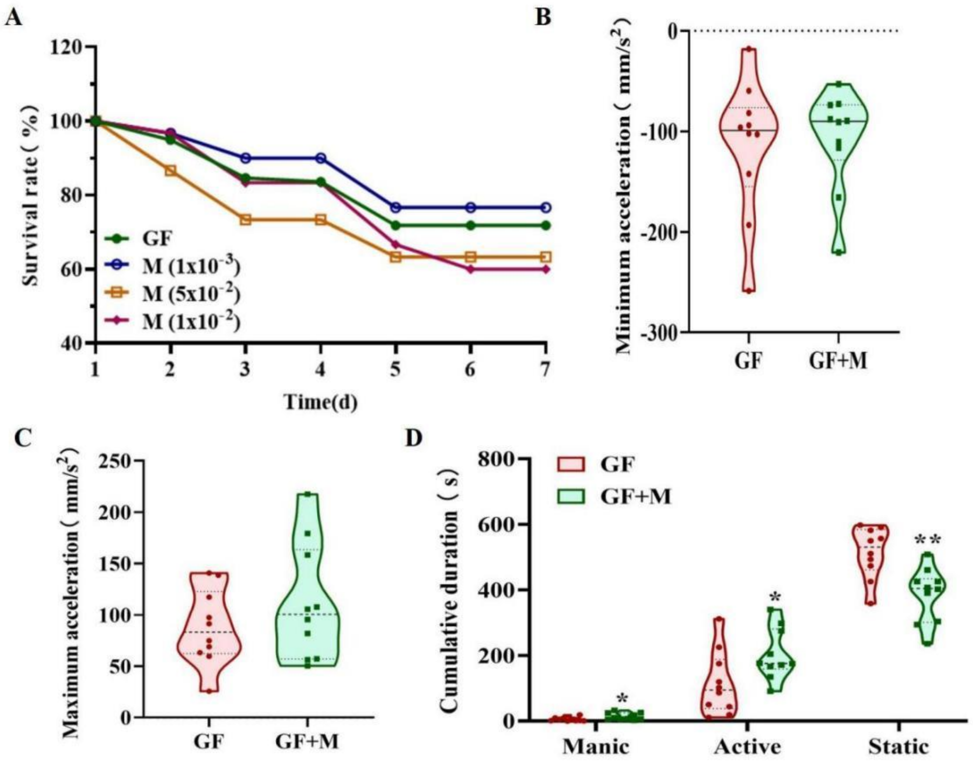


**Fig. S6. Effects of intestinal metabolites on the growth and development of GF *Oryzias latipes*.** (A) Survival rate (%) of M-exposed GF. (B–C) Minimum and maximum acceleration, including manic activity. (D) Cumulative duration of active and static states.

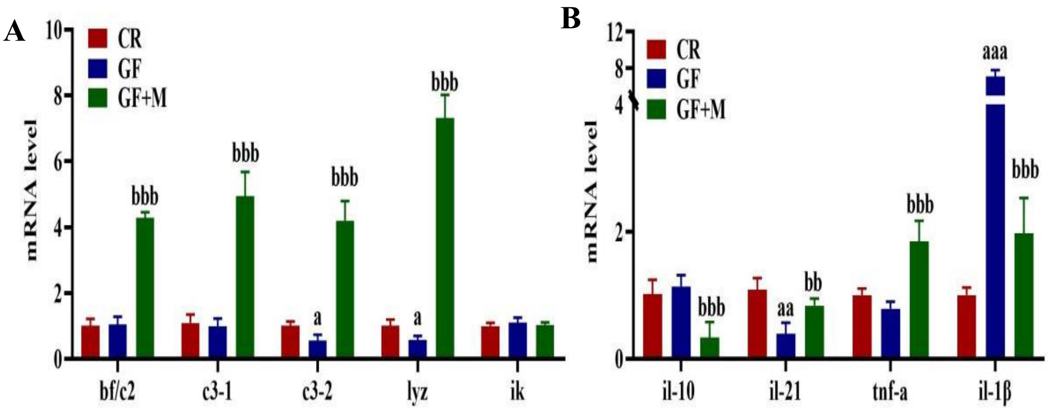


**Fig. S7. Effects of metabolites on the immune properties of GF *Oryzias latipes*.** (A–C) Expression of immune-related genes. a, aa, aaa: GF vs. CR, $P < 0.05$, $P < 0.01$, $P < 0.001$; b, bb, bbb: GF+M vs. GF, $P < 0.05$, $P < 0.01$, $P < 0.001$.

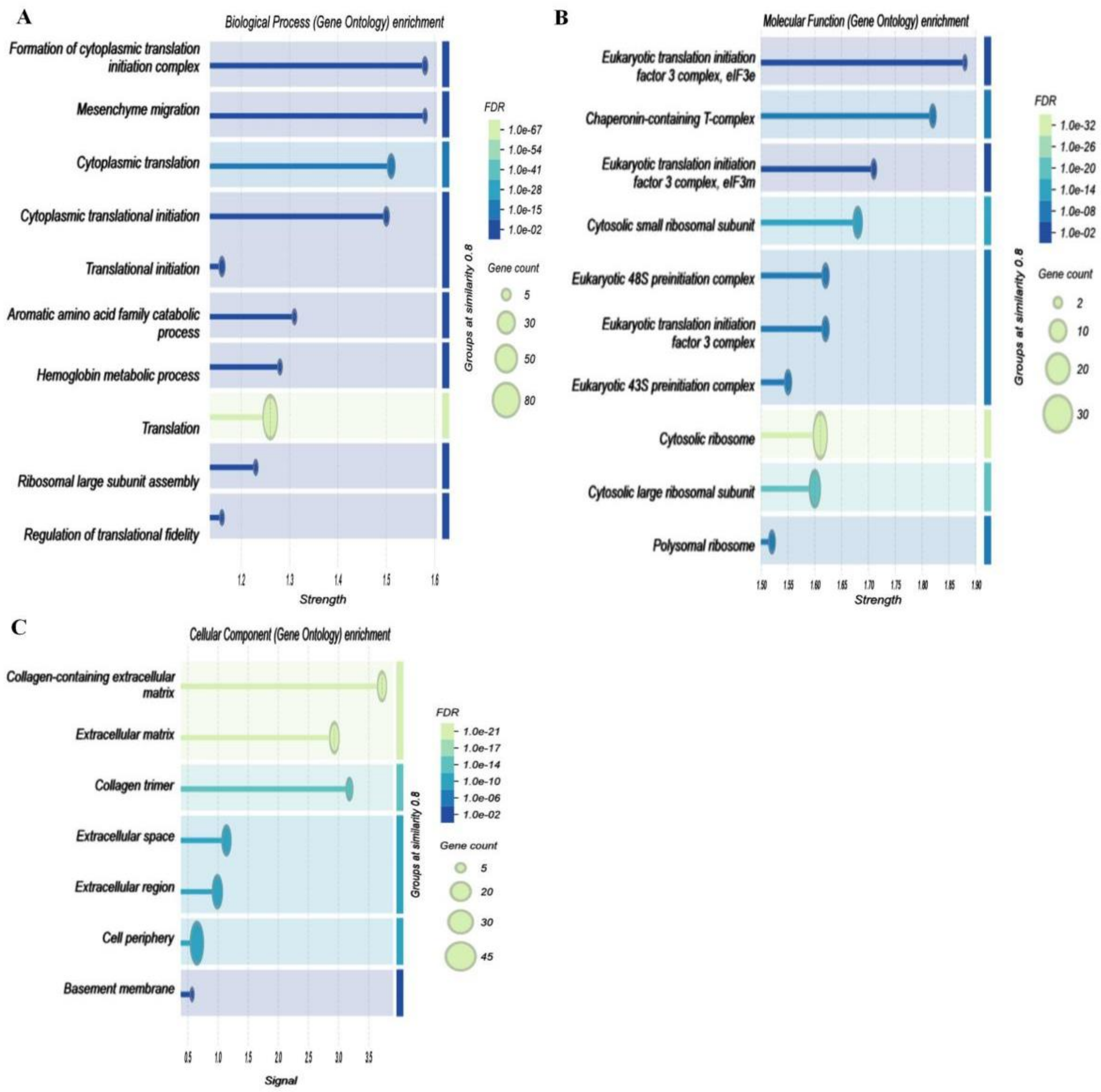


**Fig. S8. Transcriptome analysis of gut-derived metabolites in GF *Oryzias latipes*.**

(A–C) GO enrichment analysis of differentially expressed genes.

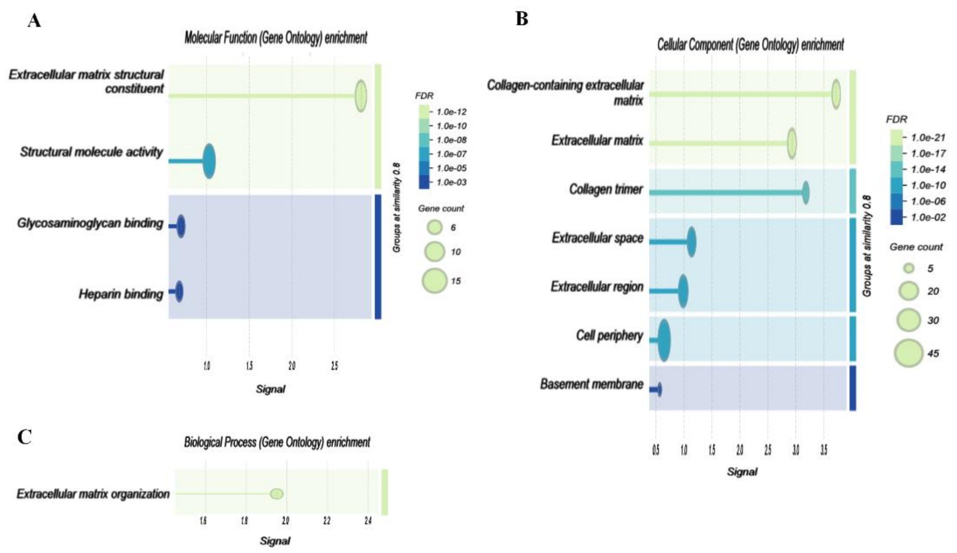


**Fig. S9. Transcriptome analysis of gut-derived metabolites in GF *Oryzias latipes*.** (A–C) GO enrichment analysis of differentially expressed genes.

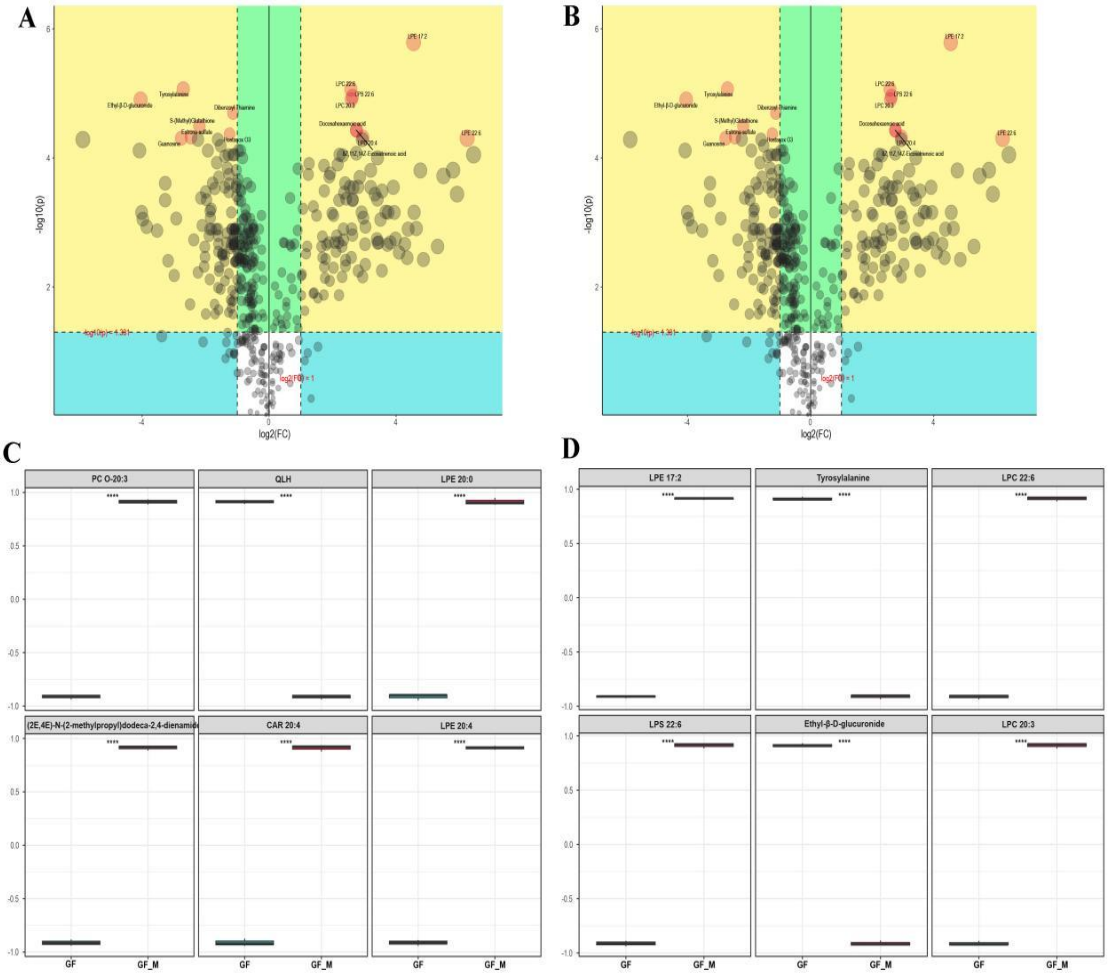


**Fig. S10. Metabolome analysis of gut-derived metabolites in GF *Oryzias latipes*.** (A–B) Volcano plots of differential metabolites in positive and negative ion modes. (C–D) Box plots of differential metabolites in positive and negative ion modes.
